# Ecological speciation promoted by divergent regulation of functional genes within African cichlid fishes

**DOI:** 10.1101/2022.01.07.475335

**Authors:** Madeleine Carruthers, Duncan E. Edgley, Andrew D. Saxon, Nestory P. Gabagambi, Asilatu Shechonge, Eric A. Miska, Richard Durbin, Jon R. Bridle, George F. Turner, Martin J. Genner

**Author notes:** Correspondence to (MC). Department of Genetics, Evolution and Environment, University College London, London, WC1E 6BT, UK.

## Abstract

Rapid ecological speciation along depth gradients has taken place independently and repeatedly in freshwater fishes. While the extent of genomic divergence between ecomorphs is often well understood, the molecular mechanisms facilitating such rapid diversification are typically unclear. In Lake Masoko, an East African crater lake, the cichlid *Astatotilapia calliptera* has diverged into shallow littoral and deep benthic ecomorphs with strikingly different jaw structures within the last 1,000 years. Using genome-wide transcriptome data from jaw tissue, we explore two major regulatory transcriptional mechanisms, expression and splicing QTL variants and examine their contribution to differential gene expression underpinning functional phenotypes. We identified 7,550 genes with significant differential expression between ecomorphs, of which 4.2% were regulated by *cis*-regulatory expression QTLs, and 6.4% were regulated by *cis*-regulatory splicing QTLs. There were also strong signals of divergent selection of differentially expressed genes that showed divergent regulation from expression, splicing or both QTL variants, including genes associated with major jaw plasticity and adaptation networks, adaptive immune system response, and oxidoreductase processes. These results suggest that transcriptome plasticity and modification have important roles during early-stage ecological speciation and demonstrate the role of regulatory-variants as important targets of selection driving ecologically-relevant divergence in gene expression that is associated with adaptive diversification.

---

Ecological opportunity has the ability to enable large-scale phenotypic diversification, potentially leading to ecological speciation among closely related species and populations^1–3^. Commonly, populations diverge in response to different selective pressures along ecological gradients, and understanding the dynamics of such organism-environment interactions is a fundamental goal in eco-evolutionary research^4, 5^. In freshwater fishes, ecological diversification along the habitat-depth gradient into contrasting trophic ecomorphs has evolved independently and repeatedly^6–11^, providing powerful natural systems to study mechanisms of ecological speciation. Here we focus on cichlid fishes, which represent valuable candidates for such investigations given their widespread ecological speciation and adaptive radiation^6, 9^. However, while their phenotypic diversity has been well-documented, and substantial insights into the genetics of population divergence in these systems have been achieved in recent years^6, 9, 12, 13^, the molecular basis and mechanisms facilitating such rapid, ecologically-driven diversification are less well understood.

The transcriptome links the genotype and phenotype and is shaped by both genetic and environmental factors^14^, offering a fundamental tool in our understanding of how environmental variation and its effects on gene expression affect local adaptation and speciation^15, 16^. Recent genome-wide studies have revealed widespread variation in gene expression among populations undergoing adaptation to ecologically divergent environments^17–22^. In addition, variation in gene expression and regulation has been shown to have a substantial heritable component upon which selection can act^23, 24^. Therefore, genetic variants regulating gene expression are likely to play a major role in facilitating adaptive diversification, and ultimately ecological speciation.

To understand how divergent gene expression and regulation might promote ecological diversification in nature, we generated whole transcriptome data from a population of cichlid fishes currently undergoing the early stages of sympatric speciation. In the East African crater lake, Lake Masoko (also known locally as Kisiba), *Astatotilapia calliptera* has diverged along a depth gradient into shallow-water “littoral” and deep-water “benthic” ecomorphs within the last 1,000 years (∼500 generations)^25^. Phenotypic and genomic divergence between these ecomorphs has been previously well-characterised^25^, revealing that substantial divergence in craniofacial morphology and feeding ecology are associated with divergence in just a small number of genetic variants. The key issue, therefore, is to understand the molecular mechanisms that generate such dramatic divergent shifts in phenotype in the absence of widespread and extensive genomic differences.

Here we explore two major heritable mechanisms of transcriptional regulation and modification, expression-quantitative trait loci (eQTLs) and splicing-QTL (sQTLs), to elucidate the molecular basis of ecological speciation. Since transcription has substantial genetic control, eQTL and sQTL mapping provides information about genetic variants with modular effects on gene expression^21, 26–31^, which are useful for understanding the genomic architecture and evolution of complex traits. Both eQTLs and sQTLs can be classified into *cis* (local) and *trans* (distant) effects. *Cis*-regulatory mutations are thought to be associated with fewer adverse effects than *trans*-regulatory mutations or protein-coding changes, and have been strongly implicated in rapid adaptation and diversification, allowing rapid trait divergence while minimising covariance with other traits (and potentially negative selective effects)^21, 30, 32^. Therefore, here we investigated variation in *cis*-acting eQTLs and sQTLs between the Lake Masoko ecomorphs to understand how transcriptional regulation facilitates divergence in functionally relevant traits during the initial stages of speciation. We predict that phenotypic diversity associated with trophic niche divergence will be facilitated by a combination of transcriptional modification from *cis*-acting expression and splicing QTLs.

In this study we first describe the extent of ecological and phenotypic diversity between the benthic and littoral *A. calliptera* ecomorphs of Lake Masoko, with specific focus on a highly adaptive cichlid trait, the lower pharyngeal jaw (LPJ), which is associated with divergence in trophic diet^33–37^. Secondly, using whole transcriptome data from LPJ tissue we investigate *cis*-acting expression and splicing QTLs to characterise the genetic architecture of regulatory variation and quantify the relative contribution of these mechanisms to divergent gene expression. Thirdly, we identify whether ecomorph-specific patterns of gene expression and regulatory variation are associated with signatures of recent selection.

## Results

### Evidence for adaptive trophic divergence in eco-morphological traits

We measured eco- morphological traits linked to trophic divergence in cichlids, namely body morphology, and LPJ morphology^34–37^ (n = 113; Supplementary Table 1), and identified differences in feeding regimes between the two ecomorphs using carbon and nitrogen stable isotope ratios. Littoral individuals had more carbon-rich diets relative to benthics (Kruskal-Wallis: chi-squared = 48.846, P = 2.131e-11; Fig. 1c), while the benthic diets were more nitrogen-rich, relative to littorals (Kruskal-Wallis: chi-squared = 17.300, P = 3.191e-05; Fig. 1c), indicating differences in trophic food source and trophic level, respectively. Bayesian stable isotope mixing models were used to determine diet compositions for benthic and littoral ecomorphs. These showed considerable differences in the contributions of dietary materials to the overall feeding regimes of the ecomorphs (Fig. 1d; see Supplementary Table 2 for further information of dietary materials). The highest diet source contributor in benthic ecomorphs was zooplankton (85.4 ± 0.079 %; mean ± sd), with a small proportion of their diet comprising of phytoplankton (5.1 ± 0.036 %; mean ± sd). The diet of littoral ecomorphs was most strongly contributed to by littoral arthropod macroinvertebrates (63.3 ± 0.346 %; mean ± sd), with additional contributions from consuming zooplankton and marginal grass (21.8 ± 0.280 % and 9.0 ± 0.067 % respectively; mean ± sd).

**Fig. 1.**
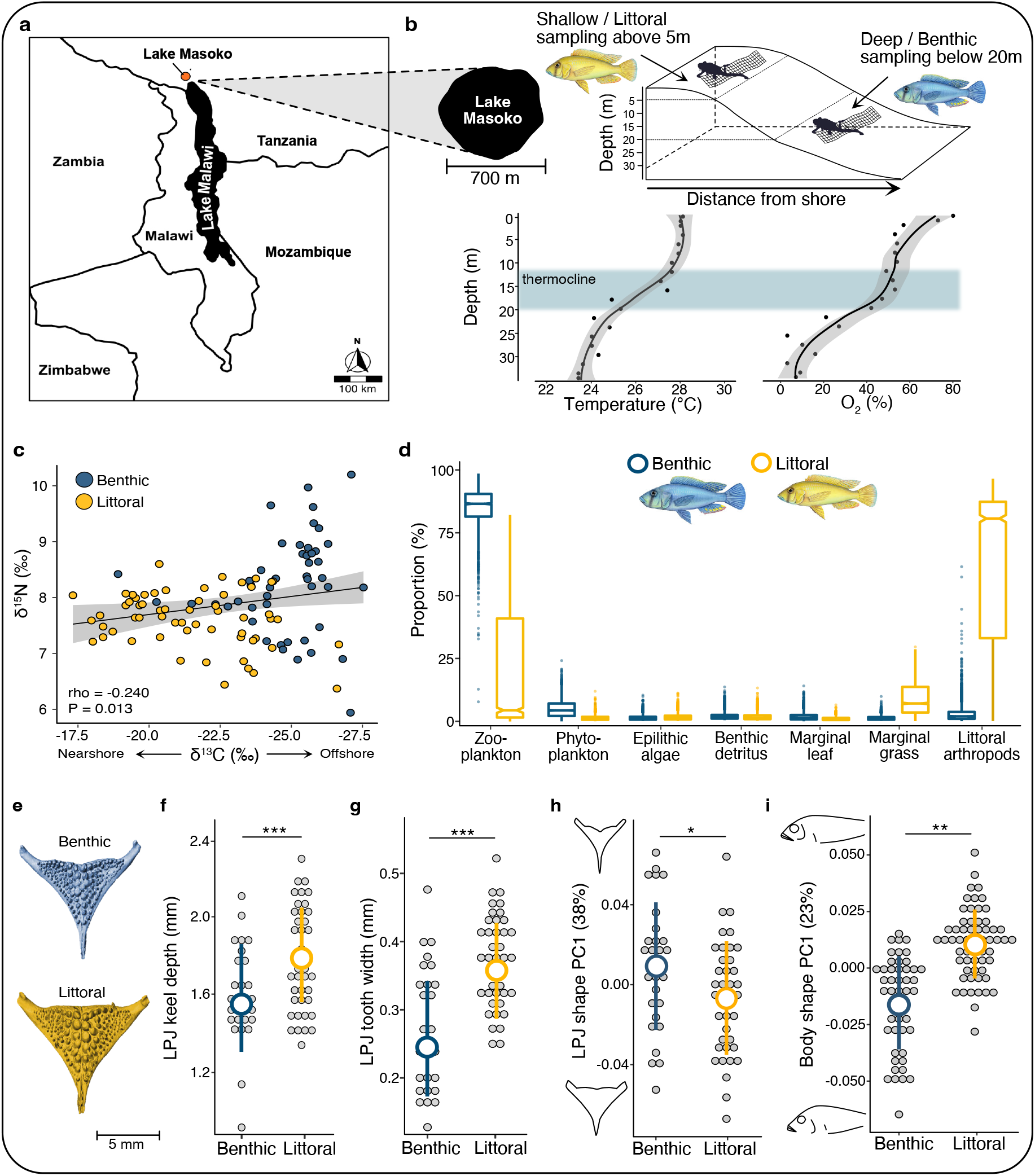
Sampling information and phenotypic divergence. **a**) The location of Lake Masoko, relative to Lake Malawi and bordering countries. **b**) Schematic of sampling approach for collection of shallow-water littoral and deep-water benthic ecomorphs of *A. calliptera* in Lake Masoko, as well as temperature and oxygen profiles by depth gradient. **c**) Association between carbon and nitrogen stable isotope signatures, demonstrating differences in trophic feeding regimes between benthic and littoral ecomorphs (n = 113). **d**) Results from Bayesian stable isotope mixing models showing proportional estimates (mean, 25% and 75% percentiles) of diet composition for benthic and littoral ecomorphs. **e)** Example of LPJ images segmented from micro-CT X-ray scans of craniofacial morphology, showing a representative example for benthic (top image) and littoral (bottom image) ecomorphs. **f-i)** Differences in LJP keel depth (n = 70), LPJ tooth width (n = 70), LPJ shape (n = 70), and body shape (n = 113), respectively, between benthic and littoral ecomorphs. Grey points represent individual samples, with mean ± sd values for each ecomorph represented by the larger coloured points and error bars (benthic = blue, littoral = yellow). LPJ shape and body shape change are shown along the first principal component axis (PC1). The proportion of variation explained is given in parentheses. LPJ and body shape outlines represent shapes at axis extremes along PC1. Number of asterisks represents the level of significance (* < 0.05, ** < 0.01, *** < 0.001).

We also found that the overall morphology of LPJs differed markedly between benthic and littoral ecomorphs (Fig. 1e). Littoral fish had significantly deeper LPJs (analysis of covariance: F_1,63_ = 12.866, P = 0.0006; Fig. 1f), and wider teeth (analysis of covariance: F_1,63_ = 28.916, P = 1.176e-06; Fig. 1g) than benthic individuals. Differences in the shape of the LPJ between ecomorphs (multivariate analysis of covariance: Wilk’s lambda_1,60_ = 0.572, P = 2.052e-06; Fig. 1h; Supplementary Fig. 1), were in the relative length and width (affects leveraging power^34, 37^), the size and position of the posterior horns (important muscle attachment sites^34, 37^), and the shape of the dentigerous area. Differences in body shape were also clear between benthic and littoral ecomorphs (multivariate analysis of covariance, Wilk’s lambda_1,58_ = 0.566, P = 2.592e-06; Fig. 1i; Supplementary Fig. 2), including body depth, head depth and length, size and position of the pectoral fin (important for manoeuvring^38–40)^, and size and position of the mouth (associated with feeding modality^41, 42^). Taken together, these results indicate a substantial and consistent shift in feeding behaviour and trophic niche specialisation between the benthic and littoral ecomorphs.

### Divergent gene expression underlies adaptive phenotypes

Genome-wide gene expression was studied using 38 whole transcriptomes from LPJ tissue (18 benthics, 20 littorals), with an average sequencing depth of 37 million reads per library (Supplementary Table 3). Cleaned reads were mapped against the *Maylandia zebra* reference genome (UMD2a, NCBI assembly: GCF_000238955.4)^43^, with a mean mapping success of 84% (Supplementary Table 3). Global expression patterns were initially explored using a Principal Component Analysis (PCA; based on the complete dataset of 19,237 genes; after filtering for low counts). The PCA demonstrated a clear separation in the overall expression profiles of benthic and littoral individuals along the primary axis of variation (PC1; Fig. 2a). A remarkably high proportion of genes (39%) were significantly differentially expressed (DE) (FDR-corrected P < 0.05; Fig. 2b; Supplementary Table 4), consistent with a high capacity for environmental plasticity^44^. Moreover, we found that both the proportion and magnitude (extent of expression divergence measured as log_2_ fold change) of DE genes upregulated in one ecomorph relative to the other was consistent (51% of DE genes upregulated in benthics, 49% in littorals; Fig. 2b). This consistent level of expression modulation across both ecomorphs suggests widespread upregulation and downregulation have facilitated phenotypic divergence in Masoko^25^.

**Fig. 2.**
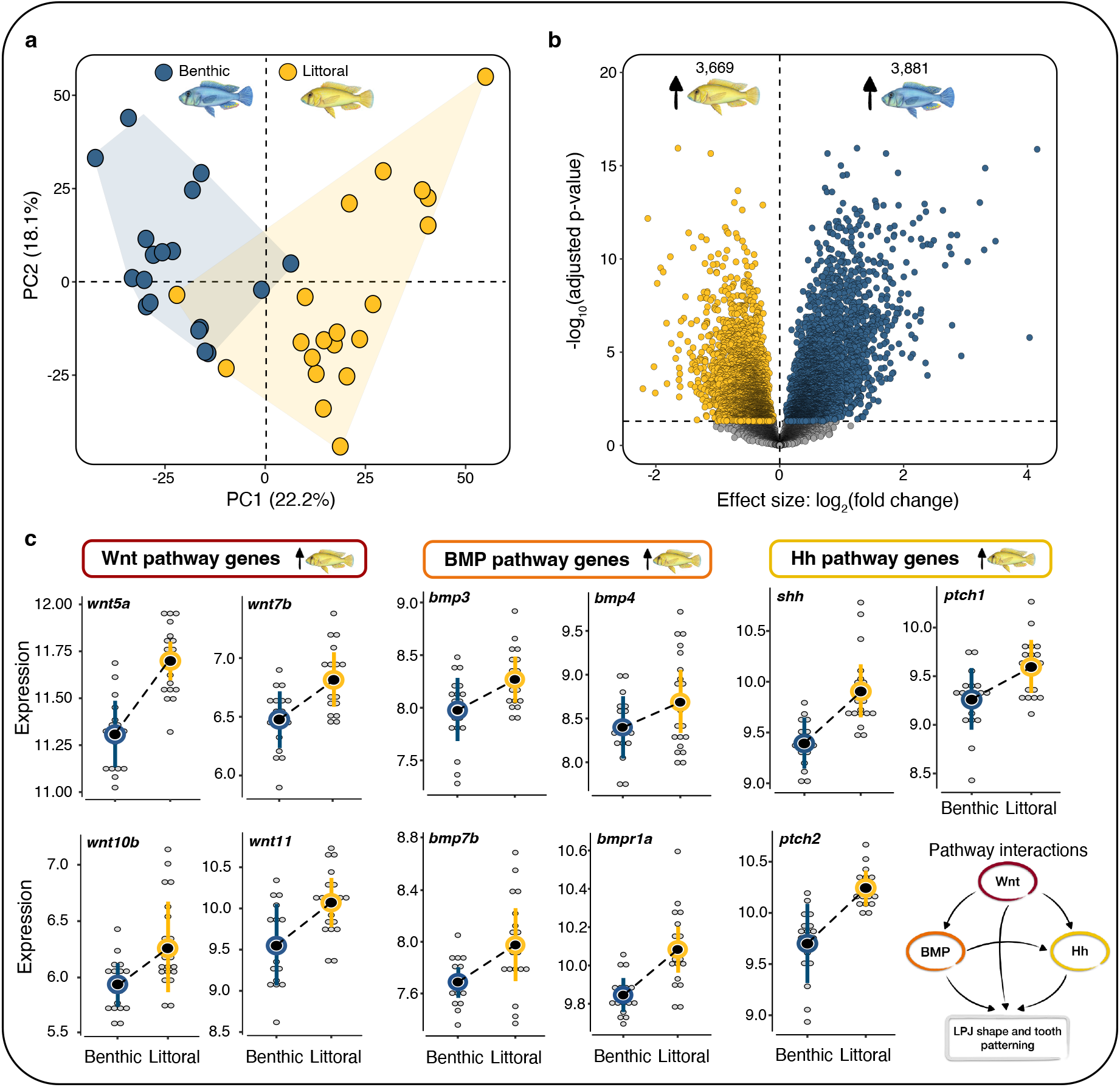
Divergent gene expression. **a**) Principal component analysis showing variation in gene expression profiles of benthic (blue; n = 18 individuals) and littoral (yellow; n = 20 individuals) ecomorphs along principal components 1 (PC1) and 2 (PC2). The percent of total variation explained by each principal component axis is given in parentheses. **b**) Volcano plot highlighting the extent of divergence in gene expression. Significant differentially expressed genes are coloured according to the expression direction (yellow = significantly upregulated in littoral, blue = significantly upregulated in benthic; FDR < 0.05). Gene expression analyses are based on the complete set of expressed genes after filtering for low counts (n = 19,237 genes). **c)** Differential expression of ‘master adaptation genes” involved in three major pathways implicated in LPJ plasticity networks and trophic adaptation (Wnt-signalling, BMP-signalling and Hh signalling). Grey points represent individual samples, with mean ± sd values for each ecomorph represented by the larger coloured points and error bars (benthic = blue, littoral = yellow). A schematic of pathway interactions during LPJ shape and tooth development is given in the bottom right.

Next, we investigated transcriptional evidence for functional differences between ecomorphs using functional enrichment analysis of Gene Ontology (GO) biological process annotations associated with DE genes. GO terms enriched in benthic ecomorphs were related to blood cell development, vascular morphogenesis and metabolic oxidation-reduction processes. This may indicate local adaptation to increased habitat depth^22, 45^ and, more specifically, to the relatively low oxygen conditions of the deep-water benthic environment in Lake Masoko, compared to shallow-water habitat (Fig. 1b and Supplementary Table 5). Interestingly, we also found significant enrichment of several immune response processes. Variation in parasite exposure, and consequently expression of immune gene networks, has been proposed as a major selective pressure and driver of adaptation and evolution in ecologically diversifying populations^46–48^. We therefore conducted an analysis of gill ectoparasites. We found that the gills of both ecomorphs were parasitised by a single species, the planktonic copepod *Lamproglena monodi*. However, as expected given their different habitats, infection rates were significantly higher in benthic (83%, n = 15 of 18) compared to littoral (36%, n = 9 of 25) individuals (GLM: Z = -2.885, P = 0.004; Supplementary Fig. 3).

GO terms enriched in littoral ecomorphs were involved in several ecologically relevant functions, including to calcium pathways, bone remodelling, muscle contraction, and neural development (Supplementary Table 5). We also found that 86% (73 out of 85) of genes previously implicated in cichlid jaw plasticity networks were differentially expressed between the Lake Masoko species pair (Supplementary Table 6). Specific genes included several bone morphogenic proteins (BMPs), *bmp3*, *bmp4*, *bmp7b*, and *bmpr1b* (Fig. 2c), which play key roles in craniofacial development^49, 50^. Moreover, *bmp4* is a known driver of craniofacial divergence during adaptive radiation in cichlid complexes^33, 51–53^, and Darwin’s finches^54^. In both cases, expression variation in *bmp4* is functionally linked to differences in foraging strategy. The consistent significant up-regulation of several Wnt (Wingless-related integration site) and Hh (Hedgehog) pathway genes in littoral ecomorphs, including *wnt5a, wnt7b, wnt10b* and *wnt11* and *shh, ptch1* and *ptch2* (Fig. 2c and Supplementary Tables 4 and 6). Given the known importance of Wnt and Hh pathways and their interactions with BMP pathways in LPJ shape and tooth patterning^51, 55–58^, these results evidence the role of divergent expression regulation in facilitating phenotypic shifts a major adaptive trait, the LPJ.

### Large effect expression and splicing QTLs regulate divergent gene expression

While gene expression can be highly plastic in its response to environmental cues, adaptive evolution of gene expression relies on a heritable genetic basis. To understand the genetic basis and mechanisms facilitating such substantial shifts in transcriptional expression profiles among ecomorphs, we conducted a genome-wide search of two major regulatory mechanisms expression and splicing QTLs. From a set of 89,718 high-confidence SNPs and indels, we located 5,722 *cis*-eQTLs that demonstrated ecomorph-specific divergent regulation of 1,001 genes (5.2% of the total expressed genes; FDR-corrected P < 0.05; Supplementary Table 7). GO analysis of *cis*-eQTL target genes revealed enrichment of several ecologically relevant GO terms, including immune response processes, DNA structural modification, cell dedifferentiation, actin cytoskeleton organisation and cilium assembly (n = 16; FDR-corrected P < 0.05; Supplementary Table 8). Cytoskeletal organisation and cilium assembly are of particular interest given that cytoskeletal signalling is recognised as a key function within LPJ plasticity networks^33, 59^, and cilia are suggested to play an important role mechanically regulated bone formation^60^ (Supplementary Table 8). We further identified 4,587 *cis*-sQTLs associated with ecomorph-specific regulation of excision ratios in one or more intron clusters in 1,422 genes (7.4% of the total expressed genes; FDR-corrected P < 0.05; Supplementary Table 9). Genes under divergent regulation from *cis*-sQTLs were enriched for a large number of GO terms, which were predominately related to immune system processes, as well as responses to bacteria and other biological stimuli (n = 97; FDR-corrected P < 0.05; Supplementary Table 10). Interestingly, we also found enrichment of terms involved in positive regulation of the ERK/MAPK (Extracellular signal-Regulated Kinase/Mitogen-Activated Protein Kinase) cascade, which is a known central regulator of tooth development in vertebrates^61–64^, as well as processes involved in regulation of cell adhesion and cell junction assembly which have been implicated in lip morphogenesis linked to foraging strategy in a Lake Tanganyika cichlid^65^.

To quantitatively assess the relationship between divergent gene expression, and *cis-* regulatory expression and splicing QTLs, we searched for overlap in the ecomorph-specific gene sets identified from each analysis (i.e., DE genes, *cis*-eQTL genes and *cis*-sQTL genes). In total we found that 4.1% of DE genes were under divergent regulation from *cis*-eQTLs, representing 31% of detected eQTLs (n = 310 genes, Hypergeometric test: P = 2.858e-07; Fig. 3a and Supplementary Table 11), 6.4% of DE genes were under divergent regulation from *cis*-sQTLs, representing 34% of detected sQTLs (n = 481 genes, Hypergeometric test: P = 1.943e-17; Fig. 3a and Supplementary Table 11), and 0.8% of DE genes were under divergent regulation from both expression and splicing QTLs (n = 63 genes; Hypergeometric test: P = 4.843e-09; Fig. 3a and Supplementary Table 11). In Lake Masoko, the benthic and littoral ecomorphs exhibit very low average genomic divergence, with only a small number of genomic regions of high divergence restricted to a few specific (‘islands’) of speciation^26^. Therefore, to determine whether a similar pattern was observed for transcriptional divergence associated with overlapping gene sets we examined the genomic distribution of the 728 genes that were identified as having differential expression patterns mediated by one or more *cis*- regulatory QTL variants (total *cis*-eQTLs = 1,290, total *cis*-sQTLs = 2,539). We found that regions associated with extensive transcriptional divergence were more or less evenly distributed across the genome and were not conserved to specific genomic regions (Fig. 3b). This suggests that large regions of the transcriptome are undergoing divergence through *cis*- regulatory elements.

**Fig. 3.**
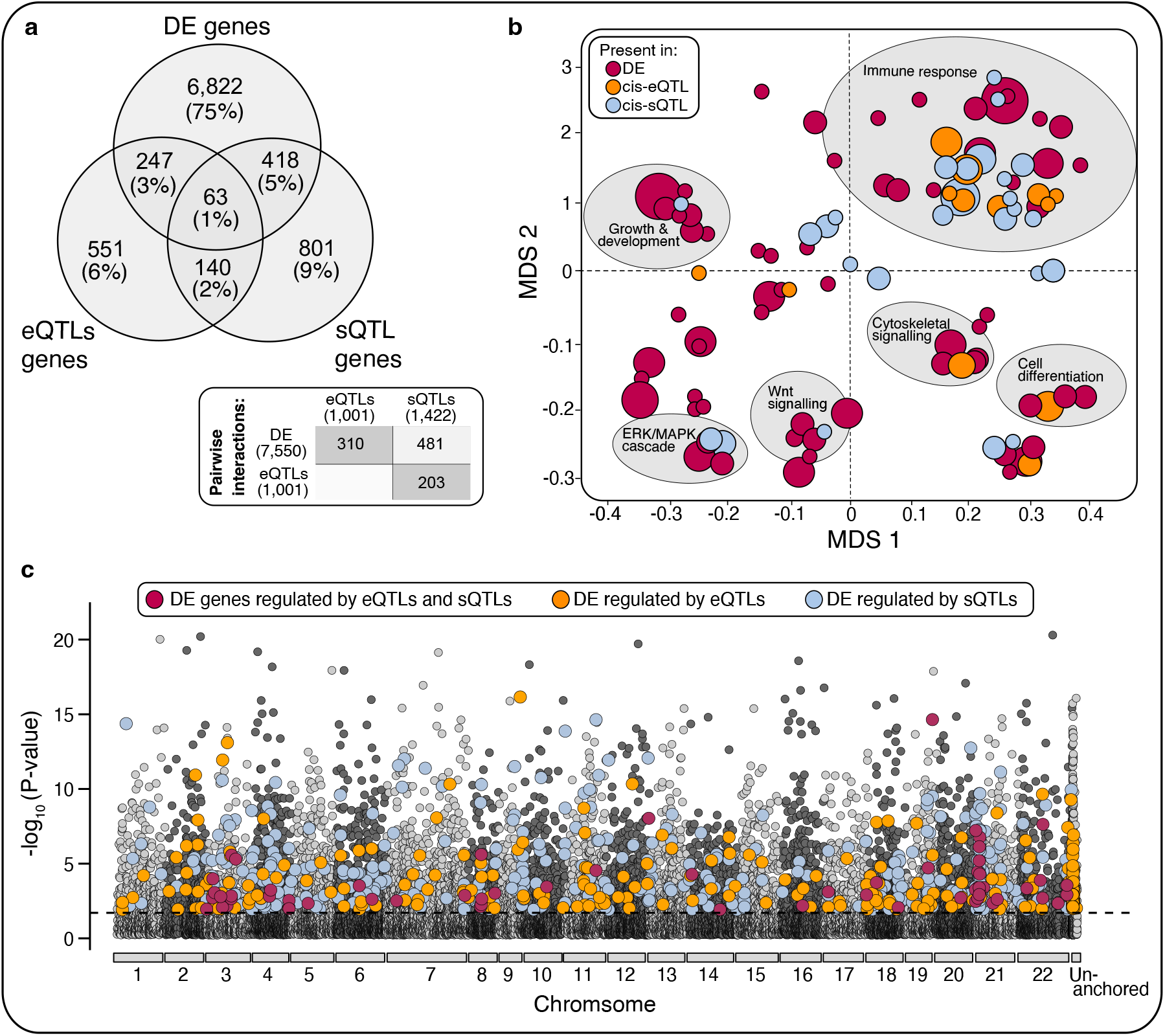
Cis-regulatory elements associated with expression variation between ecomorphs. a) Venn diagram showing the number and proportion of genes that overlap between differential expression (DE), *cis*-regulatory splicing QTLs (*cis*-sQTL) and *cis*-regulatory expression QTLs (cis-eQTL) analyses. **b)** Multi-dimensional scaling (MDS) plot showing enriched biological process GO terms identified for significant gene sets. Red circles represent enriched terms for DE genes, orange circles represent enriched terms for *cis*-eQTL genes, and blue circles represent enriched terms for cis-sQTL genes (FDR < 0.05). GO terms were clustered based on semantic similarity of functional classifications. The size of the circle corresponds to the number of genes associated with each GO term cluster. Descriptions are given for GO cluster functions that were shared across all three datasets (DE, *cis*-eQTL and *cis*-sQTL genes). **c)** Manhattan plot showing the genomic distribution of differentially expressed (DE) genes that were associated with divergent regulation from expression or splicing QTLs (according to their position in *M. zebra* reference genome). DE genes regulated by *cis*-eQTL variants are represented by orange circles, DE genes regulated by *cis*-sQTL variants are represented by blue circles and DE genes regulated by both *cis*-eQTL and *cis*-sQTL variants are represented by red circles. Chromosomes are highlighted by alternating colours and un-anchored scaffolds are located at the right end of the x-axis. The y-axis relates to the magnitude of expression divergence in individual genes, given as -log_10_ transformed FDR-corrected P-values generated from the global differential expression analysis (based on the total set of 19,237 expressed genes). The black dashed line indicates the 5% FDR threshold.

In line with these results, we identified a diverse range of functions that were enriched for DE that were regulated by expression, splicing or both QTL mechanisms (Fig. 3c and Supplementary Table 12). We found that DE genes under divergent *cis*-regulation from either expression or splicing QTLs, or both, were enriched for several GO terms with potential implications in LPJ adaptation and trophic diet specialisation, including Wnt and cytoskeletal signalling processes which have established roles in LPJ plasticity networks^33, 59, 66^, as well as ERK/MAPK pathway regulation, metalloendopeptidase activity, cell adhesion and cell junction assembly, all of which have recognised roles in tooth development and regeneration^61–64, 67^. Of specific interest, we identified two genes, *bmp7b* and *ptch2*, that have been repeatedly linked to cichlid jaw plasticity and adaptation, and more specifically tooth patterning and regeneration^56, 58, 66, 68–71^, that were associated with divergent *cis*-eQTL genotypes (Fig. 4). For both genes, *bmp7b* and *ptch2*, we found that the level of expression was higher in homozygous individuals for the major allele, compared to both heterozygous and minor allele homozygous individuals. In both cases, reduced expression and minor allele genotypes were more typically associated with benthic individuals, which show narrower, papilliform LPJ phenotypes, with significantly smaller teeth than littoral individuals (Fig. 1 and Fig. 4). These results provide the first evidence of an at least partial genetic basis to this highly plastic and complex adaptive trait during the early stages of sympatric speciation in Lake Masoko.

**Fig. 4.**
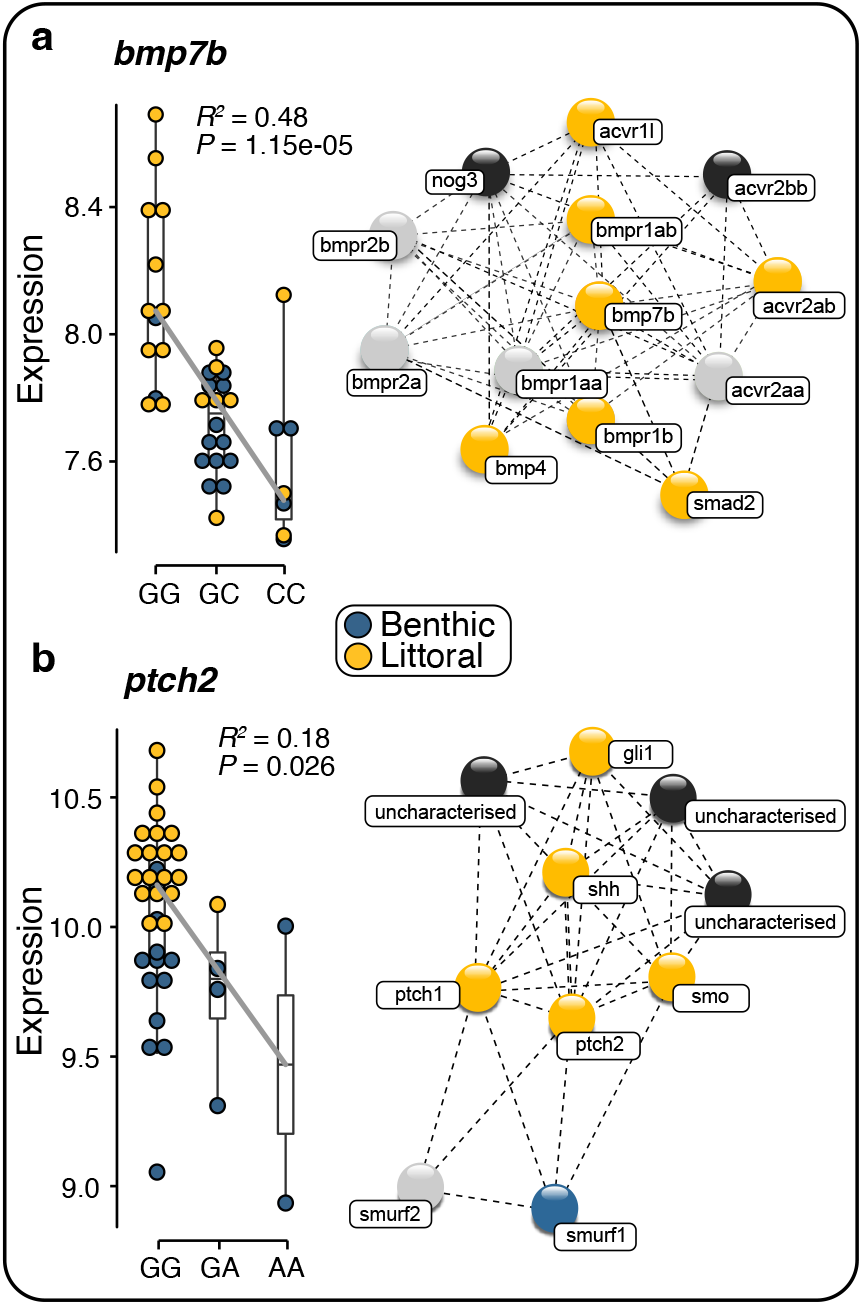
Divergent regulation in candidate genes underlying LPJ trait divergence. Associations between *cis*-eQTL genotypes and the level of expression (normalised counts) in two candidate genes involved in LPJ adaptation pathways, **a)** *bmp7b* (Bone morphogenic protein 7b) within the BMP-signalling pathway and **b)** *ptch2* (Patched 2) within the Hh-signalling pathway. Gene abundance per individual, per genotype is shown as blue circles for benthic individuals and yellow circles for littoral individuals. Significance of genotype- specific specific expression in both genes of interest was tested using individual linear models. The grey line corresponds to the linear model fitted to the data and associated statistics (coefficient of determination: R^2^ and *p*-value: P) are given in each panel. Schematics of gene network interactions for both candidate genes are shown next to the corresponding genotype plots. Gene interactions were deduced based on available data for *Danio rerio* and *Maylandia zebra* in STRING (https://string-db.org). Blue gene nodes represent significant upregulation in the benthic ecomorph, yellow nodes represent significant upregulation in the littoral ecomorph, grey nodes represent genes that were recovered but showed no significant differential expression or regulation between ecomorphs, and black nodes represent genes that were not recovered in our dataset.

Of further interest, we found that shared GO terms identified across all three gene sets (DE genes, *cis*-eQTL genes and *cis*-sQTL genes) were involved in a diverse range of immune system processes, including divergent regulation of the major histocompatibility complex (MHC), which has a widely recognised role during local adaptation^46–48^. Notably, divergence in MHC alleles has been linked to host-parasite coevolutionary dynamics during adaptive trophic divergence in haplochromine cichlids within the Lake Malawi radiation^72^. We also identified expression and splicing QTL mediated upregulation of several processes involved in oxidoreductase activity, angiogenesis, integrin-mediated signalling, and blood coagulation cascades (Supplementary Tables 8, 10 and 12), which were almost exclusively associated with up-regulation in benthic individuals. Specific genes included histone deacetylase 6 (*hdac6*), hepatic and glial cell adhesion molecule a (*hepacama*) and prostaglandin G/H synthase 2 (*ptgs2*) which are directly involved in the activation of the hypoxia-inducible factor pathway^73, 74^ (Supplementary Fig. 4). All three genes are significantly upregulated in benthic individuals further indicating a potential role of divergent expression regulation in promoting adaptation to increased depth and reduced oxygen conditions in the deep-benthic habitat. Taken together, these results highlight both expression and splicing *cis*-acting QTLs as significant regulators of differential expression underlying several ecologically relevant phenotypes and provides an invaluable insight into the molecular basis of transcriptional modifications facilitating trophic niche adaptation.

### Evidence for divergent selection facilitated by large-effect *cis*-eQTLs

To further investigate the functional genomic basis and potential role of *cis*-regulatory variants in transcriptome evolution and ecological speciation we applied two approaches. First, we looked at the association between ecomorph-specific expression and splicing QTLs and genome-wide differentiation (*F*_ST_, estimated using non-overlapping 10kb sliding window averages to avoid bias from any single highly differentiated SNPs within a given window). We found that *cis-*regulatory eQTL and sQTL variants associated with differential gene expression were significantly more likely to be located in genomic regions with higher genetic differentiation between ecomorphs (eQTL-DE: *F*_ST_ = 0.032 ± 0.0034; sQTL-DE: *F*_ST_ = 0.034 ± 0.0032; mean ± se), compared to the genome wide average (*F*_ST_ = 0.019 ± 0.0002; mean ± se; Student’s t-test_eQTL-DE_: t _(148.34)_ = -3.896, P = 0.0002; Student’s t-test_sQTL-DE_: t _(260.78)_ = -4.654, P = 5.176e-06). These results support the hypothesis that divergent natural selection is acting on *cis* mechanisms that regulate ecomorph-specific shifts in gene expression and transcriptional modification in Lake Masoko *A. calliptera*. More specifically, given that *cis* regulatory changes accumulate preferentially over time^29, 31, 32, 75–77^, and that *cis*- regulatory divergence is shown to increase linearly with divergence time^78^, the high levels of *cis*-regulatory divergence observed here in such a young species pair (less than 1,000 years old) suggests that *cis*-mechanisms are a major driver of radiation within this cichlid lineage.

Second, we determined whether patterns of ecomorph-specific divergence in gene expression, and expression and splicing QTLs were associated with genetic signatures of selection. We performed genome scans using two complementary haplotype-based statistics, xpEHH and xp-nSL to search for evidence of recent selective sweeps^79^. To determine whether genes under selection were associated with hard or soft sweeps we used a combination of H2/H1 and H12 statistics, following the approach developed by Garud et al. ^80^. Surprisingly, given the very recent divergence of this species pair, we found evidence of recent selection in 563 SNPs (from a set of 89,718 high-confidence transcriptome SNPs), which were associated with 128 genes (Fig. 5). Consistent with early-stage speciation, and selection on standing genetic variation^81–83^, we found that soft sweeps formed the dominant mode of adaptation in Lake Masoko *A. calliptera*, with 65% of SNPs under selection associated with evidence of soft sweeps, and 35% associated with hard sweeps (Supplementary Table 13). Genes associated with signatures of selection were enriched for a total of 25 GO terms, including epithelial cell development and morphogenesis, apical constriction, transferrin transport, and regulation of digestive system processes - specifically lipid absorption and oxidoreductase activities (FDR-corrected p-value < 0.05; Supplementary Table 14).

**Fig. 5.**
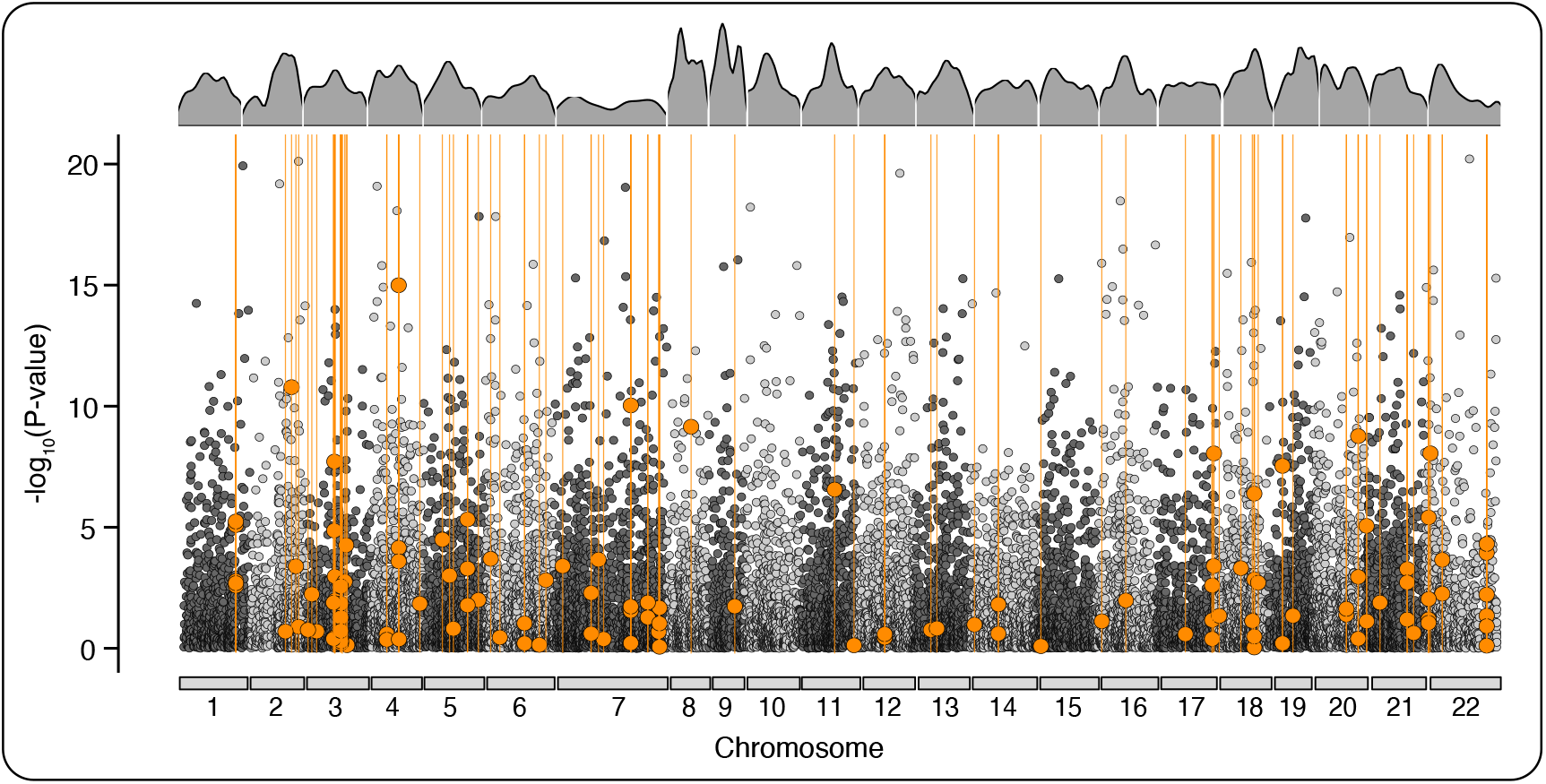
Signatures of selection. Genomic distribution of genes associated with signatures of positive selection and the associated magnitude of expression divergence, given the -log_10_ transformed FDR-corrected P-values generated from the global differential expression analysis across all anchored chromosomes (based on 19,237 expressed genes). Genes showing evidence of significant divergent selection between benthic and littoral ecomorphs are highlighted with orange points, and regions with high H12 selection scores illustrated by the orange lines. The gene density per chromosome is depicted along the top section of the plot.

We found considerable overlap between genes under selection and genes that were differentially expressed (62 out of 128; Hypergeometric test: P = 0.013), genes under divergent regulation from *cis*-eQTLs (13 out of 128; Hypergeometric test: P = 0.006), and genes under divergent regulation from *cis*-sQTLs (30 out of 128; Hypergeometric test: P = 2.679e-09) (Supplementary Fig. 5 and Supplementary Table 11). Moreover, we found three (out of 128) genes were shared across all four analyses (Hypergeometric test: P > 0.05), which were identified as *iqgap2* (IQ motif-containing GTPase-activating protein 2), *scrn3* (Secernin 3) and an uncharacterised serine-type endopeptidase protein (LOC101485919).

Interestingly, *iqgap2* has been implicated in Wnt-pathway regulation, and more specifically in tooth development in mice^84, 85^. *Scrn3* is putatively involved in blood glucose regulating, and serine-type endopeptidase proteins are often functionally linked to regulation of blood coagulation^86^, both of which are common adaptive responses to hypoxia^87, 88^. Taken together, these results provide direct evidence of positive selection on regulatory transcriptional mechanisms associated with trophic diet specialisation and occupation of habitats with different oxygen concentrations. They also suggest that regulatory variants underpinning transcriptional diversity are major facilitator of rapid ecological diversification.

## Discussion

In this study we simultaneously investigated changes in two *cis*-regulatory mechanisms, expression and splicing QTLs, and their relative contribution to divergent gene expression patterns underpinning ecological speciation in cichlids. We found that substantial transcriptional modification facilitates ecological diversification, despite an absence of fixed genomic differences and ongoing gene flow^25^. We also assessed several novel aspects of eco-morphological divergence within this species pair, including a detailed analysis of LPJ morphology (Figs. 1e-1h, and Supplementary Fig. 1). We found that the observed direction and extent of LPJ divergence was comparable to divergence patterns in species pairs in older, more established cichlid radiations^34, 35, 37, 41^. We also report the first evidence of niche-specific differences in parasite load, and divergent expression regulation of major adaptive immune response networks between ecomorphs in Lake Masoko (Supplementary Fig. 3 and Supplementary Tables 11 and 12), which has emerged recently as important driver of host speciation in freshwater fishes^72, 89–94^. The observed divergence in multiple ecomorphological traits (Fig. 1), coupled with genomic evidence of widespread recent selection, is consistent with a scenario of multifarious selection acting on numerous axes of the phenotype simultaneously.

We suggest that substantial shifts in the transcriptional landscape have allowed the rapid evolution of extensive phenotypic diversity despite a relatively low level of overall genomic differentiation between these ecomorphs. Divergent expression was identified for 24.2% of genes genome-wide (Fig. 2b), which is markedly higher than has been reported previously in comparable studies of within-lake ecomorph pairs in natural systems (1.6% in Arctic charr^22, 95^, 9.7% in Lake whitefish^96^, and 11.4% in European whitefish^96^). We further show that the ecomorph-specific gene expression patterns observed directly underpin ecologically relevant phenotypes (Supplementary Tables 5). In particular, several genes we identified have been previously highlighted by common garden diet-manipulation experiments as being directly involved in LPJ plasticity and feeding efficiency associated with differing trophic diets^33, 59^ (e.g., BMP, Wnt, and Hh signalling pathways; Fig. 2c and Supplementary Table 6), suggesting that transcriptionally-mediated adaptative diversification occurs through the refinement (and potential genetic assimilation) of plastic response to variable environments in nature. Furthermore, given that the substantial shifts in gene expression underpinning such phenotypic diversity are controlled by a relatively small number of fixed genetic variants in Lake Masoko^25^, it seems likely that pre-existing environmental plasticity has shaped the transcriptional landscape in our diversifying focal species. Such a pattern supports the ‘plasticity-first’ evolution model of adaptive divergence, where plastic phenotypic responses coupled with the evolution of habitat preference (with increasingly consistent genotype- habitat associations) lead to divergence at loci showing environment-dependent expression.

Eventually such plasticity is lost as adaptive divergence (and habitat fidelity) continues, and flexible gene expression is replaced by fixed differences in both the genome and phenotype^97, 98^.

Although widespread transcriptional plasticity has the capacity to facilitate initial phenotypic shifts towards different environments, to contribute to adaptive evolution divergent gene expression and regulation need to have a heritable genetic basis^99^. Remarkably, given the young age of our focal species pair (initial divergence estimated between 500-1,000 years ago^25^), we found that a significant proportion of differentially expressed genes were associated with changes in large-effect *cis*-regulatory variants, evidencing a genetic basis to some of the phenotypic and transcriptional diversity observed. We found that 728 (10%) of the genes exhibiting divergent expression regimes between benthic and littoral ecomorphs were regulated by at least one expression or splicing *cis*-regulatory QTL (Fig. 3). Given that *cis* regulatory changes accumulate preferentially over time^29, 31, 32, 75–77^, and that *cis*-regulatory divergence is shown to increase linearly with divergence time^78^, the high levels of *cis*- regulatory divergence observed here in such a young species pair suggests that *cis*- mechanisms are a major driver of radiation within this cichlid lineage. Moreover, we found that both expression and splicing *cis*-regulatory variants were associated with more highly differentiated genomic regions, implicating *cis* mechanisms as important targets of natural selection during the early stages of ecological speciation.

We further show that target genes of expression and splicing QTL are associated with several ecologically relevant functions, including the divergent regulation of BMP, Hh, and Wnt signalling pathways which have widely recognised roles in shaping craniofacial phenotypes during trophic niche specialisation and adaptive radiations in cichlids^51, 55–58^. We also identified *cis*-QTL mediated divergent regulation of ERK/MAPK cascades between the benthic and littoral Masoko ecomorphs (Supplementary Table 12). ERK/MAPK signalling plays a major in tooth and skeletal development and has been previously shown to be an important trophic modulator in mice^62, 64^, and more recently as an important driver of trophic modulatory in oral and pharyngeal jaws in cichlids^63^. Additionally, we identify two genes as potential candidate underpinning LPJ divergence between the benthic and littoral ecomorphs in Lake Masoko, *bmp7b* (BMP signalling) and *ptch2* (Hh signalling). Both genes play central roles in tooth development, patterning and regeneration, with differential expression regulation of *bmp7b* and *ptch2* linked to differences in feeding behaviour and niche diversification in mice^68^. More specifically, *bmp7b* and *ptch2* are recognised ‘master adaptation genes’ and have been repeatedly implicated in LPJ adaptation and plasticity networks^33, 59, 66^ (Fig. 2 and 4, and Supplementary Table 5). Therefore, we suggest that the observed transcriptional shifts in expression and regulation of these genes play a fundamental role in shaping adaptive LPJ shape and tooth phenotypes during trophic diversification in our focal species. This also is consistent with a heritable genetic basis to these complex phenotypes upon which selection can act. In support of this prediction, we found evidence of recent selection on several genes involved in epithelial cell morphogenesis and apical constriction, including the gene *iqgap2* which showed evidence of divergent regulation both through expression and splicing QTLs, and has been functionally linked to tooth development in humans^84, 85^.

Interestingly, we also found significant *cis*-mediated modification and selection of several genes involved in adaptive response immune system networks, including the major histocompatibility complex, and we also found that these were predominantly associated with adaptation towards the deep-benthic environment (Supplementary Tables 12, 13 and 14).

Divergence in MHC alleles has been shown to drive host-parasite coevolutionary dynamics within the Lake Malawi cichlid radiation^72^. Given that benthic individuals exhibited an increased parasite load compared to littoral individuals in the current study, the observed transcriptional shifts in MHC expression and regulation could indicate that similar host-parasite co-evolutionary dynamics are ongoing in Lake Masoko. Additionally, divergence in MHC complexes have also been linked to female mate preference in three-spine sticklebacks^47, 94, 101^, meaning that such ecologically-driven changes are likely to also contribute to speciation by promoting assortative mating and reproductive isolation.

In support of the role of environmentally sensitive genomic regions shaping ongoing adaptive divergence, we found evidence of *cis*-mediated regulation and positive selection in genes involved in the activation and regulation of hypoxia-inducible factor signalling pathways and anticoagulation cascades (Supplementary Fig. 4 and Supplementary Tables 12, 13 and 14).

Formation of new blood cells has been well-documented in mammals as a response to low oxygen conditions^102–104^. Therefore, we suggest that occupation of the deep-water, near hypoxic benthic niche in Lake Masoko is facilitated by a combination of transcriptional regulation from expression and splicing and *cis*-regulatory mechanisms.

Overall, our results reinforce the role of gene expression in adaptive evolution. In particular, we show that substantial divergent modification of gene expression and transcriptional regulatory mechanisms underpin adaptive eco-morphological traits during early stages of sympatric speciation. These findings provide evidence that the vast transcriptional landscape acts as a rich substrate for innovation, allowing rapid diversification in resource use where resources become spatially or reliably distinct. These data suggest that an accumulation of environmentally induced transcriptional changes might promote genomic predisposition in lineages such as cichlids towards divergent adaptation and ecological speciation.

Furthermore, given that our focal species and study system, *Astatotilapia calliptera* from Lake Masoko, are part of the broader Lake Malawi haplochromine cichlid species flock, we suggest that our results have importance for understanding the molecular mechanisms facilitating explosive diversification in cichlids, and perhaps more broadly for large-scale adaptive radiation across a wider range of vertebrate taxa.

## Methods

### Sample collection

*Astatotilapia calliptera* were sampled from Lake Masoko, in October 2019 (Fig. 1a). Fish were collected using monofilament blocknets and SCUBA at two habitat depths: above 5m for littoral ecomorphs and below 20m for benthic ecomorphs (Fig. 1b).

Benthic fish were transferred to holding containers within the lake and brought to the surface gradually over a duration of three days to allow for decompression. All fish were held in aerated containers during transportation from the field to the Tanzania Fisheries Research Institute in Kyela. A total of 113 fish were sampled, 65 from the shallow-littoral zone and 48 from the deep-benthic zone (Supplementary Table 1). Fish were euthanised with MS-222 (Sigma-Aldrich), following an approved procedure. Immediately after euthanising, fish were photographed, and standard length was measured to the nearest millimetre. Lower pharyngeal jaws (LPJs) were dissected from a subset of 38 individuals (20 littoral, 18 benthic) and stored in RNAlater. Tissues for RNA extractions were collected within five minutes of euthanasia. All individuals were preserved in ethanol and stored at -20 C for a minimum of three days before shipment to the UK. All further processing was carried out at the University of Bristol, UK.

Scientific fish collections from Lake Masoko were carried out under research permits issued by the Tanzania Commission for Science and Technology (permit number: 2019-549-NA- 2019-357).

### Stable isotope analysis

To infer differences in feeding behaviour and trophic niche specialisation between ecomorphs, we performed stable isotope analysis on white muscle tissue samples from our focal fish (n = 113) and variety of dietary materials (n = 21). White muscle tissue was dissected from the left side of the ethanol-preserved fish, posterior to the operculum, above the lateral line to determine isotope signatures for carbon (δ^13^C) and nitrogen (δ^15^C). Dietary materials were represented by zooplankton, phytoplankton, macrophytes, littoral arthropods, epilithic algae and benthic detritus (Supplementary Table 2). All samples (muscle tissue and dietary materials) were oven dried at 60°C overnight. Stable isotope analyses were carried at Iso-Analytical, Crewe UK. Isotope ratios were identified using Elemental Analysis - Isotope Ratio Mass Spectrometry with a Europa Scientific 20-20 IRMS. Carbon and nitrogen isotope compositions were calibrated relative to VPDB and AIR, respectively, using IA-R042 (NBS-1577B: powdered bovine liver) as reference material.

Duplicate analyses were performed on 20% of samples as a control, and instrumental accuracy was monitored using multiple reference materials (IA-R042: powered bovine liver, IA-R045/IA-R005: a mixture of ammonium sulphate and beet sugar, and IA-R046/IA-R006: a mixture of ammonium sulphate and beet sugar).

To assess overall differences in trophic in diet between benthic and littoral ecomorphs we performed separate Kruskal-Wallis tests for carbon and nitrogen stable isotope ratios, using standard functions in R *v.*4.0.2^105^. The relative proportions of potential dietary materials consumed by benthic and littoral individuals were determined using the R-package simmr^106^, which applies a Bayesian Markov Chain Monte Carlo (MCMC) approach to estimate the composition of sources (i.e., dietary materials) that contribute to the resulting isotope signatures identified for your targets (i.e., focal species groups). The simmr MCMC model was run for 100,000 iterations with a burn rate of 1,000.

### Morphological analysis

To quantify phenotypic variation between ecomorphs we measured several morphological traits, including head and body shape, LPJ shape, LPJ keel depth, LPJ tooth width and LPJ tooth length (Supplementary Fig. 6).

Morphological variation in LPJs was quantified using 3D reconstructions of LPJs using micro-computed tomography (micro-CT) scans of whole specimens (n = 70; 40 littoral, 30 benthic). A Nikon XTH225ST micro-CT scanning system was used to generate the images. Each scan used 2,750 projections and included between eight and 10 individuals. Scan resolution was determined by the size of region of interest (i.e., the head and upper section of the body), and as such was determined by the overall body size of each individual. The average voxel size across scans was approximately 23 μm. Image stacks were exported into *VG Studio v.*3.0 (Volume Graphics GmbH, 2016) and 3D models were reconstructed for each individual. We used *Avizo v.*8.0 (Hillsboro, OR) to isolate the LPJ from the rest of the skeleton and capture 2D images of the dorsal and superior-lateral perspectives. LPJ shape was analysed using a landmark-based geometric morphometric approach, as described above. To accurately capture the shape of curved edges along the LPJ, we used a combination of standing-homologous and sliding semi-landmarks (Supplementary Fig. 6). Standing landmark positions followed Muschick et al. ^34^. Images were scaled and digitised with *tpsDig2 v*.2.30^107^. Procrustes superimposition was conducted in *MorphoJ v*.1.07a^108^ to standardise landmark configuration and remove any unwanted variation related to the size, position and orientation of the fish. Principal component analysis (PCA) was used to identify the major axes of variation in LPJ shape between littoral and benthic ecomorphs. Linear measurements for LPJ keel depth, tooth width and tooth length were also collected using *tpsDig2*. For tooth width and length, the posterior three teeth immediately to the right side of the suture were measured (to the nearest 0.01mm) and averaged (Supplementary Fig. 6).

Head and body shape was analysed from 2D images using a landmark-based geometric morphometric approach (n = 113). All individuals were photographed in a standard orientation with the head pointing to the left. Homologous landmark positions were selected based on Malinsky et al. ^25^ (Supplementary Fig. 6). Images were calibrated to scale, and landmarks were digitised using *tpsDig2 v*.2.30^107^. Procrustes superimposition was conducted in *MorphoJ* following the same methods described for LPJ shape analyses. Principal component analysis (PCA) was used to identify the major axes of variation in head and body shape between littoral and benthic ecomorphs.

Analysis of covariance models were used to test for significant ecomorph-specific differences in all morphological traits, using body size (standard length), sex, and their interaction as covariates. Non-significant terms were excluded from final models (body size and sex were non-significant in all models).

### RNA extraction, sequencing and mapping

Total RNA was extracted from LPJ tissue using RNeasy Mini Kits (Qiagen), following the manufacturer’s instructions with minor modifications. Modifications included a two-step homogenisation, using a TissueLyser LT (Qiagen) followed by QIAshredder (Qiagen) column homogenisation. We also performed two on-column washes with 80% ethanol immediately prior to the final RNA elution to remove any residual buffer. RNA quantity and quality were assessed using the Qubit 2.0 fluorometer (Life Technologies) with BR Assay kits and a 2100 Bioanalyser platform (Agilent), respectively. High quality RNA was achieved, with A260/280 ratios between 1.9 and 2.1 and RNA Integrity Numbers above 7. RNA-seq libraries were prepared and sequenced at the Bristol Genomics Facility (University of Bristol). Separate cDNA libraries were prepared for each individual. Libraries were prepared using TruSeq Stranded mRNA Sample Preparation Kits (Illumina), in combination with a Poly-A selection step. Sequencing was performed on an Illumina NextSeq 500 platform (Illumina), using 75 bp paired-end sequencing at a sequencing depth of approximately 30 million reads per individual.

Raw reads were processed before mapping. Adapters were removed with *Scythe v*.0.9944 BETA (http://github.com/vsbuffalo/scythe/) and low-quality reads were removed with *Trimmomatic v.*0.36^109^. *FastQC* v. 0.11.8^110^ was used to assess the quality of reads before and after pre-processing. Cleaned reads were then mapped to the *Maylandia zebra* reference genome (UMD2a, NCBI assembly: GCF_000238955.4)^43^ with *STAR v.*2.7.1a^111^. Mapping was performed using the two-pass mode in STAR in order to identify splice junctions and allow subsequent analysis of splice sites. HTSeq *v.*0.11.1^112^ was used to quantify gene expression and generate read count tables.

### Differential gene expression analysis

Data were filtered to remove genes with less than 10 reads across 50% of samples, and the filtered counts were log_10_ transformed using the *rlog* function in the R-package *DESeq2 v.*1.28.1^113^. We performed a PCA on the *rlog*-transformed read counts, using the R-package *PCAmethods v.*1.81.0^114^, to examine the overall gene expression profiles. To determine the extent of variation in gene expression between littoral and benthic ecomorphs, we conducted a differential expression (DE) analysis using negative binomial distribution models in *DESeq2*. All *p*-values were adjusted for multiple testing using Benjamini and Hochberg correction^115^ (FDR < 0.05).

### SNP genotyping and effect predictions

Single nucleotide polymorphisms (SNPs) and indels were called using the physical mapping information from the *STAR* two-pass alignments of the RNAseq data, for all samples. Before calling SNPs, duplicates were marked and removed using *Picard*-*tools v*.2.20.0 (https://broadinstitute.github.io/picard/).

SNPs were called using *Freebayes v*.0.9.21^116^, specifying a minimum coverage of three reads per sample to process a site and a minimum of two reads per sample to consider an alternative allele. Using *VCFtools v*.0.1.16^117^, we filtered the resulting SNP dataset to retain only biallelic SNPs, with a phred quality above 30, genotype quality above 30, a minor allele frequency of 5%, were present in at least 90% of all individuals, and passed a Hardy-Weinberg disequilibrium threshold of *p*-value < 0.01. Further filtering was performed using the *vcffilter* program within *vcflib*^118^, specifying an allele depth balance 0.30 and 0.70. This resulted in a final set of 89,718 high-confidence, gene-associated SNPs.

### *cis*-eQTL analysis

We conducted eQTL mapping analysis to identify genetic variants putatively underlying divergent gene expression patterns between littoral and benthic ecomorphs. We focused solely on *cis*-eQTLs because we lack the substantial power required for *trans* identification in with the current dataset size (n = 38). *Cis*-eQTLs were identified with the R package *MatrixEQTL v*.2.3^119^. We applied the linear model approach within the *MatrixEQTL* package to test for gene expression variation in response to genotype variation. Genotypes for eQTL analysis were generated from the complete set of 89,718 gene-associated SNPs (called from RNAseq data) using the 012-recoding function in *VCFtools*, and normalised expression counts for the set of 19,237 expressed genes were formed the expression phenotype dataset in the model. As suggested by Shabalin ^119^ we checked our expression dataset for outliers prior to running the model using the provided code (no outliers were detected). To control for population structure in our *cis-*eQTL analysis, we performed a PCA on the complete set of 89,718 SNPs using *Plink2 v*.2.00a.2.3^120^. Only the first principal component (PC1) was associated with ecomorph-specific genotype variation (analysis of covariance: F_1,36_ = 48.694, P = 3.507e-08; Supplementary Fig. 7; P > 0.2 for all remaining PCs) and was used as a covariate in the *MatrixEQTL* model (gene expression ∼ genotype + PC1_genotype_ + sex). We defined *cis*- eQTLs as being within 1 Mb of the transcription start site of their target gene. All *p-*values were corrected for multiple testing (FDR < 0.05), implemented directly by the *MatrixEQTL algorithm*. Quality control of the test model was assessed using histogram and quantile-quantile plots of the total inferred *cis*-eQTLs for the tested dataset, implemented directly within *MatrixEQTL*, indicating inflation of p-values on the right side of the plot only (Supplementary Fig. 8). To determine whether the observed inflation in extreme p-values was an effect of population structure or other spurious effects, we performed permutation testing of the model with simulated genotype data. We performed 50 genotype simulations (retaining population flags for individuals) to break genotype-expression associations. Quantile-quantile plots and number of significant *cis*-eQTLs detected were compared between our test model and the simulated data models to assess accuracy of our results. The inflated p-values observed in our test model were not reflected by the simulated data models, and the number of significant *cis-* eQTLs detected for simulated data models ranged between 1 and 19 (compared to 5,722 significant *cis*-eQTLs in our true data), indicating that our results are reflective of true genotype-gene associations and not a result of spurious effects (Supplementary Fig. 9 and Supplementary Table 15).

### *cis*-sQTL analysis

To generate splicing phenotypes for our sQTL mapping we estimated intron excision ratios with *LeafCutter v*.0.2.9^121^ following the authors’ recommended pipeline. Briefly, *LeafCutter* uses bam file alignments from STAR two-pass mapping as input and generates ratios of reads supporting each alternatively excised intron. Introns with ratios < 0.001 and used in less than 40% of individuals were removed resulting in a final set of 73,311 alternatively excised intron clusters across 13,295 genes. The remaining ratio values were log_2_ transformed and underwent intron-wise quantile normalisation and were used as the splicing phenotype for QTL mapping. *MatrixEQTL* was used to perform *cis*-sQTL mapping following the same approach described for our *cis*-eQTL mapping analysis. To test for variation in intron excision in response to genotype variation we implemented the linear model approach in *MatrixEQTL*, using the set of 75,311 alternatively excised intron clusters as the phenotype data input and the set of 87,918 SNPs as the genotype input data. Sex was included as a fixed effect and population structure was accounted for by including the first PC from the PCA analysis on genotypes as a covariate (intron excision ratios ∼ genotype + PC1_genotype_ + sex). We defined *cis*-sQTLs as being within 1 Mb of their target intron cluster. All *p-*values were corrected for multiple testing (FDR < 0.05), implemented directly by the *MatrixEQTL* algorithm. Quality of the test model was assessed using histogram and quantile-quantile plots of the total inferred *cis*-sQTLs for the tested dataset, implemented directly within *MatrixEQTL*. Again, inflation of p-values was observed on the right side of the quantile-quantile plot (Supplementary Fig. 10), therefore permutation testing was performed using simulated genotype datasets, using the same approach applied to our *cis*-eQTL dataset. Quantile-quantile plots and number of significant *cis*-sQTLs detected by permutation models indicated that the results of our test model are reflective of true genotype-phenotype associations with zero to nine significant *cis*-sQTLs detected for simulated data models (Supplementary Fig. 11 and Supplementary Table 16).

### Association between gene expression divergence and genomic differentiation

To determine whether differentially expressed genes under divergent regulatory control from *cis*- acting expression and splicing QTLs play a role in transcriptome evolution, we examined whether they overlap with genomic regions of high genetic differentiation (*F*_ST_). Weir and Cockerman’s *F*_ST_ was calculated for the complete set of 89,718 SNPs identified from the LPJ RNAseq dataset. We used non-overlapping 10kb sliding window averages to avoid bias from any single highly differentiated SNPs within a given window, implemented in *VCFtools*. Genome-wide window-based *F*_ST_ values were tested against averaged *F*_ST_ values for the set of genes undergoing ecomorph-specific divergent regulation from splicing and *cis*-regulation by performing a Student’s t-test using R built-in functions^105^.

### Signatures of selection

We performed phasing and missing genotype imputation on the complete set of SNPs (n = 89,718) with *Beagle v*.4.1^122^. Haplotype matrices and physical maps were generated for each chromosome separately using *Plink2 v*.2.00a.2.3^120^. Genome- wide signatures of selection were then identified using a combination of two complementary haplotype-based statistics to scan for evidence of recent positive selection across all anchored chromosomes (n = 22 chromosomes, n = 69,390 SNPs). Specifically, we estimated xpEHH (cross-population extended haplotype homozygosity) and xp-nSL (cross-population number of Segregating sites by Length) statistics using *selscan v*.1.3.0+^79^. To identify candidate SNPs under selection we calculated the top and bottom 1% quantiles for normalised scores of both xpEHH and xp-nSL statistics (calculated using the *norm* function within Selscan), and SNPs shared across both approaches were considered as being under selection. Finally, to infer whether SNP-linked genes under selection were associated with hard or soft sweeps we used a combination of H2/H1 and H12 statistics, using the H12_H2H1.py script from Garud et al.^80^. H2/H1 and H12 statistics were calculated for each anchored chromosome separately using overlapping windows of 25 single nucleotide variants, with a step size of 1, and allowing zero false positive single nucleotide variants per window. High H12 scores were determined using the H12 critical value, i.e., greater than the genome-wide median of H12. Where high H12 scores were associated with high H2/H1 ratios (greater than 0.05), selection was considered to be influenced by soft sweeps. SNPs under selection were linked with a given gene if they were within 1 Mb upstream of the transcription start site.

### Co-distribution of key regulatory mechanisms and patterns of divergent expression and selection

We investigated the genomic co-distribution of all gene sets of interest (i.e., DE genes, *cis*-eQTL genes, *cis*-sQTL genes, and genes linked with SNPs putatively under selection). We tested performed hypergeometric tests with the R-package *SuperExactTest* to test the probability that genes were shared more often than expected by chance. Tests were performed using 10,000 simulations and specifying the total set of 19,237 expressed genes as the background gene set to inform the significance of overlap between gene sets of interest. Significance was assessed using Fisher’s exact tests with an FDR correction for multiple testing (FDR < 0.05).

### Functional enrichment analysis

Gene ontology (GO) annotation was performed using the R package *topGO v.*2.41.0^123^, based on *Danio rerio* UniProt annotations. Functional enrichment analyses of GO biological process terms were identified using the *parentchild* algorithm in *topGO*. Lists of genes of interest identified by DE, *cis*-eQTL, *cis*-sQTL and selection analyses were individually compared against the reference set of 19,237 annotated genes in the *M. zebra* genome. *P*-values for GO terms were obtained using a Fisher’s exact test with an FDR correction for multiple testing (FDR < 0.05). Significant GO terms were clustered based on semantic similarity using the *ViSEAGO v.*1.3.16 clustering algorithm^124^ and visualised using multidimensional scaling (MDS) plots. Hypergeometric tests were used to assess whether significant GO clusters associated with each gene set of interest were shared more than expected by chance, following the methods outlined above.

## Acknowledgments

Many thanks are due Joseph Masore from the Tanzania Fisheries Research Institute in Kyela for his assistance during field work, and to Charles Malela and Thomas Katerso for their enormous effort and help during sample collection. We thank Zacharia J. Mwampwani for his fundamental logistical support during fieldwork. We thank Thomas Davies and Elizabeth Martin-Silverstone for their advice and assistance in generating micro-CT scans. We thank Stella Jones and Alex A. Perez for their contribution to the collection of parasite data. We thank Guillermo P. Gonzalez for his comments and insight. Finally, we thank Julie Johnson for creating the illustrations of our focal species. Sampling permission was issued by the Tanzania Commission for Science and Technology, permit number 2019-549-NA-2019-357. Research was funded by a NERC grant NE/S001794/1 to MJG, GFT and JRB. Ethical approval for this project was awarded by the Animal Services Unit at the University of Bristol (UIN 19054).

## Author contributions

MJG, GFT and JRB designed the project, with support from EAM and RD. MC, DEE, NPG and AS conducted and facilitated fieldwork. MC and ADS generated the phenotypic data. MC conducted the phenotypic analyses. MC generated, performed bioinformatics and all analyses of the transcriptome data, and compiled and analysed the comparative analyses. MC produced the figures. MC and MJG wrote the manuscript with contributions from GFT and JRB. All authors contributed to interpreting the results and revising the manuscript.

## Data availability

Raw sequence data and all phenotypic and transcriptomic datasets presented in this paper, and any required code will be submitted to the Environmental Information Data Centre (EIDC), hosted by the UK Centre for Ecology and Hydrology (UKCEH), on acceptance.

