## Supplementary figures 1-11 for "Ecological speciation promoted by divergent regulation of functional genes within African cichlid fishes"

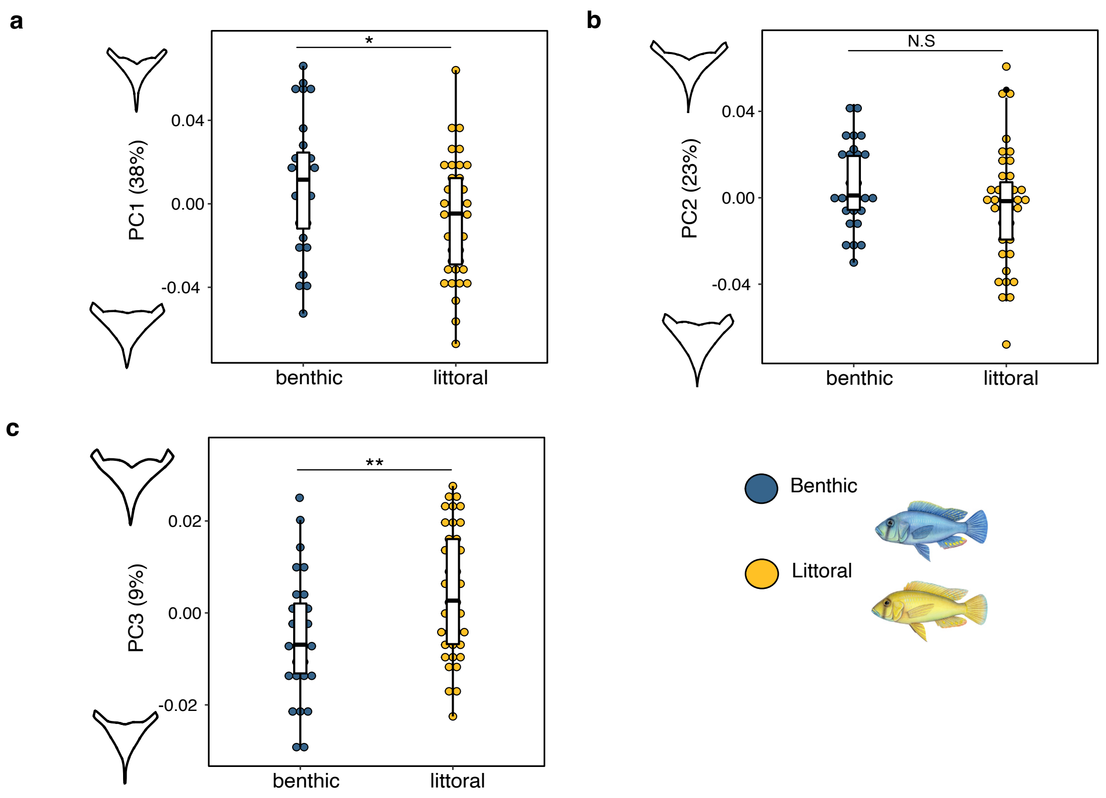


**Supplementary Figure 1 | Major axes of LPJ shape change. a)** LPJ shape change along PC1. **b)** LPJ shape change along PC2. **c)** LPJ shape change along PC3. The percent of expression variation explained by each axis is given in parentheses. Outline shapes represent axis extremes for all PCs. Asterisks denote significant differences between benthic (blue) and littoral (yellow) ecomorphs. Number of asterisks represents the level of significance. N.S represents non-significant differences between ecotypes. N = 70.

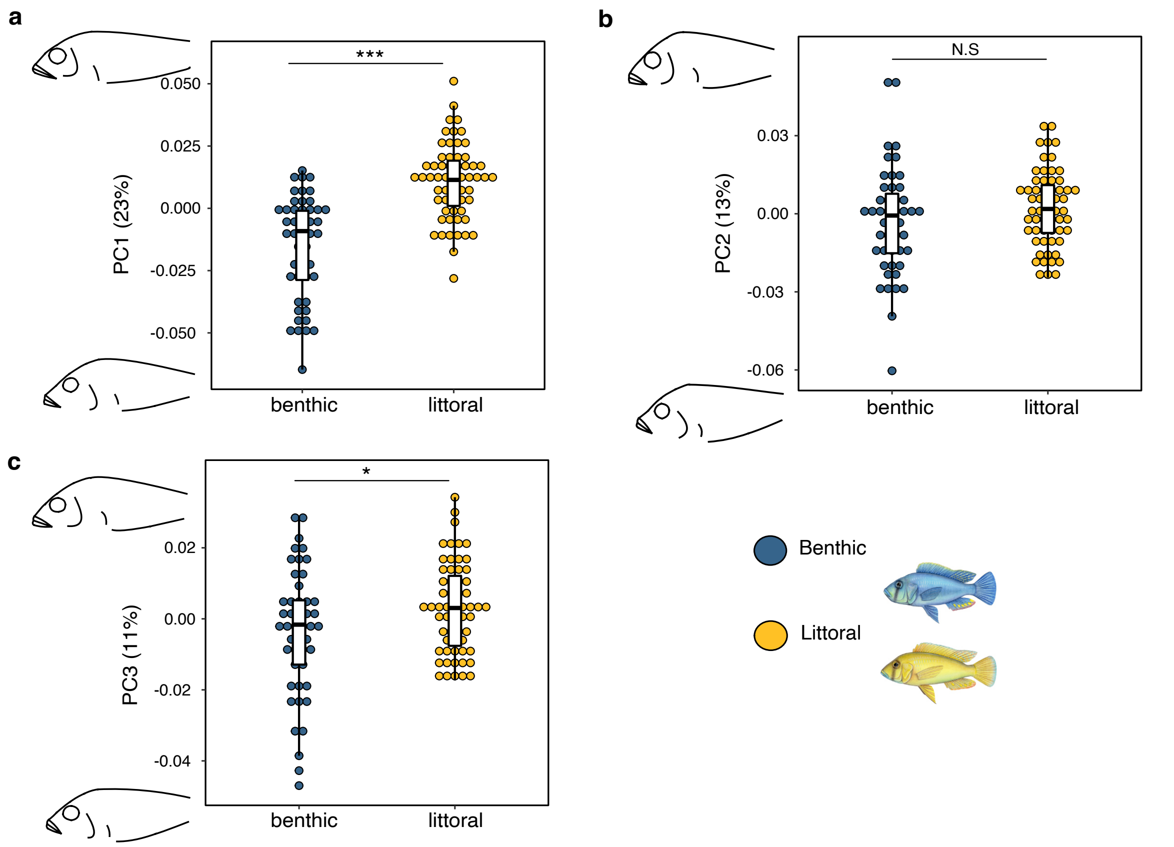


**Supplementary Figure 2 | Major axes of body shape change. a)** Body shape changes along PC1. **b)** Body shape changes along PC2. **c)** Body shape changes along PC3. The percent of expression variation explained by each axis is given in parentheses. Outline shapes represent axis extremes for all PCs. Asterisks denote significant differences between benthic (blue) and littoral (yellow) ecomorphs. Number of asterisks represents the level of significance. N.S represents non-significant differences between ecomorphs. N = 113.

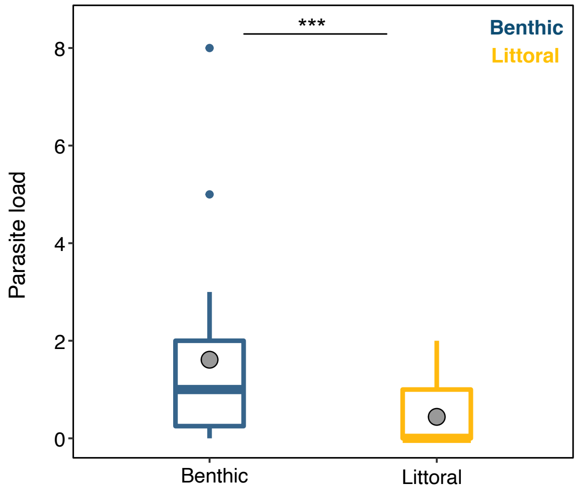


**Supplementary Figure 3 | Parasite load differences.** Difference in parasite loads of the gill ectoparasite species, *Lamproglena monodi* identified for benthic (blue) and littoral (yellow) ecomorphs. Grey points represent them mean values for each ecomorphs. Asterisks denote significant differences between benthic and littoral ecomorphs. Number of asterisks represents the level of significance. N = 38.

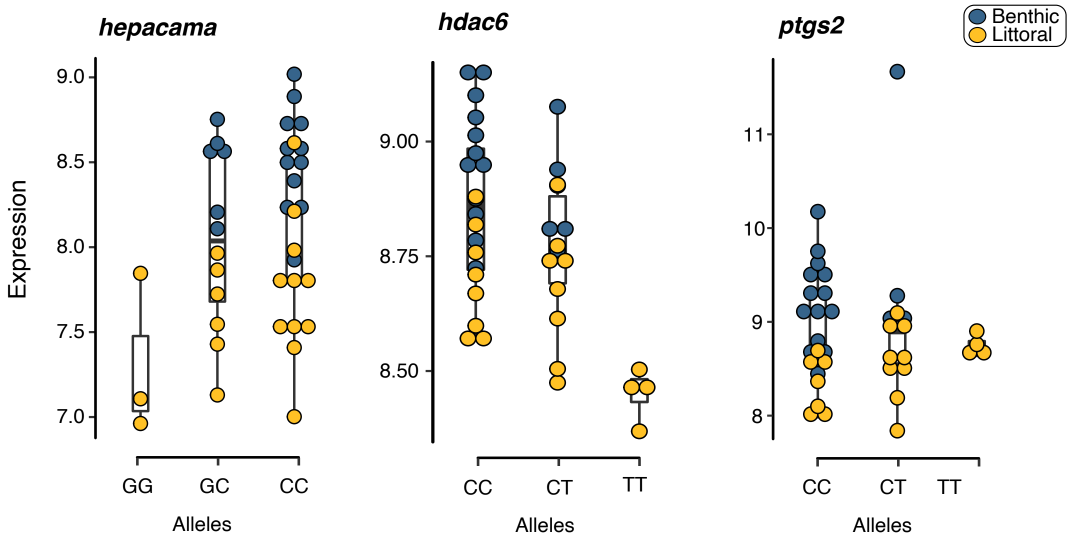


**Supplementary Figure 4 | Genes involved in adaptive response to hypoxia.** Genotype-based expression of three candidate genes involved activation of the hypoxia inducible factor pathway during hypoxia adaptation response. All three genes were identified as being under significant DE and *cis*-QTL regulation, as well as under selection. The y-axis shows normalised expression of each gene for both ecomorphs (littoral in yellow, benthic in blue). Genotypes are given on the x-axis.

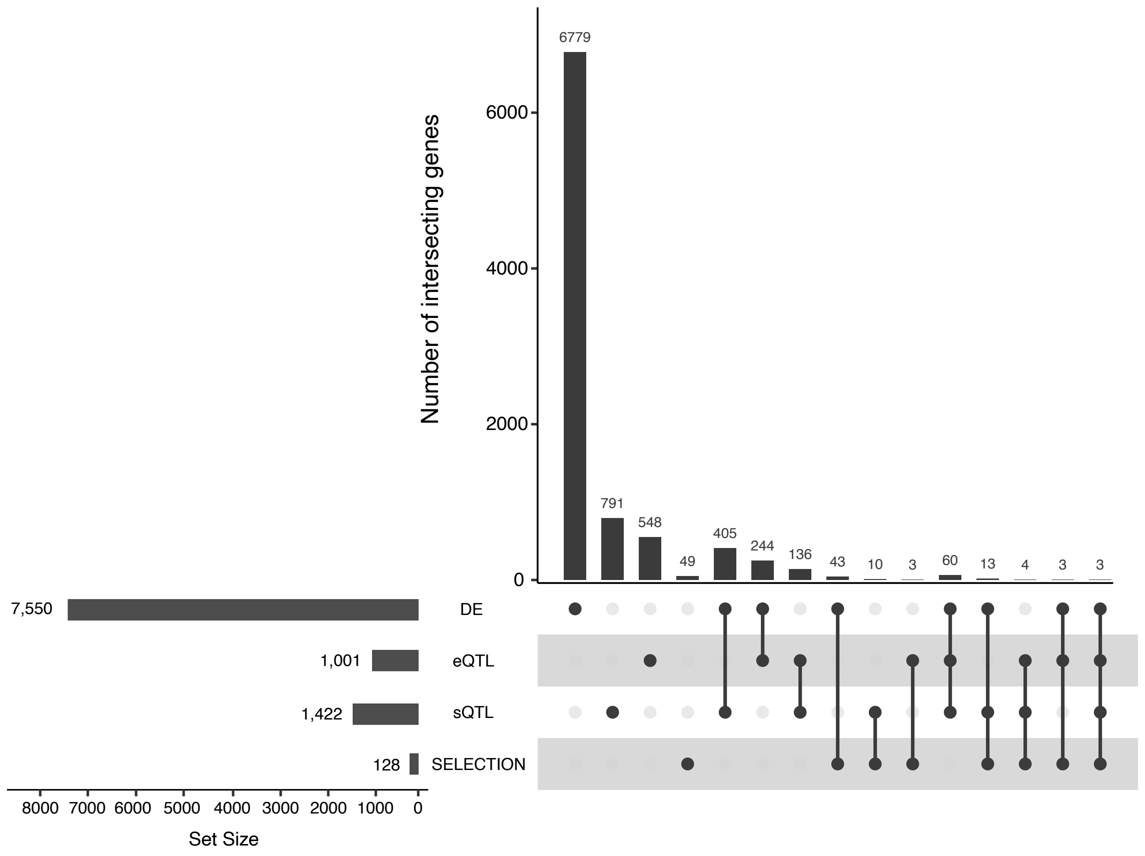


**Supplementary Figure 5 | Frequency of shared genes between transcriptional analyses.** Different analyses are indicated by the dots for DE (differentially expressed genes, n = 7,550), eQTL (*cis* expression QTL regulated genes; n= 1,001), eQTL (*cis* splicing QTL regulated genes; n= 1,422), and SELECTION (genes under selection; n = 128). Individual dots represent the number of genes unique to a given analysis. Linked dots represent the number of shared genes across analyses.

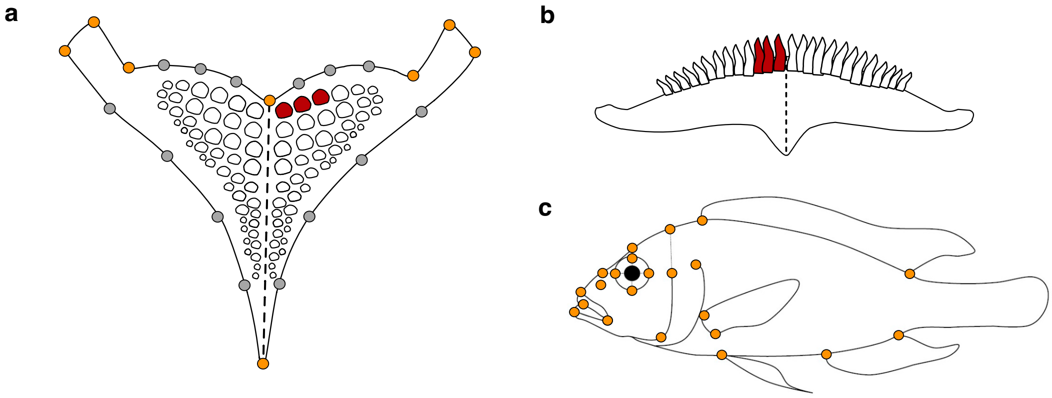


**Supplementary Figure 6 | Schematic of morphological measurements for LPJ and body shape. a)** Dorsal view of the LPJ. Landmark scheme used for geometric morphometric analysis of LPJ shape. Orange points represent standing landmark positions and grey points represent sliding semi-landmarks; total of 22 landmarks. LPJ tooth width measurements were collected from the first three posterior teeth immediately to the right of the suture line and are highlighted in red. **b)** Posterior view of the LPJ. Keel depth was measured following the suture line (dashed line). The first three teeth to the right (from a dorsal perspective) were used for tooth length measurements and are highlighted in red. **c)** Landmark scheme used for geometric morphometric analysis of body shape; total of 22 landmarks.

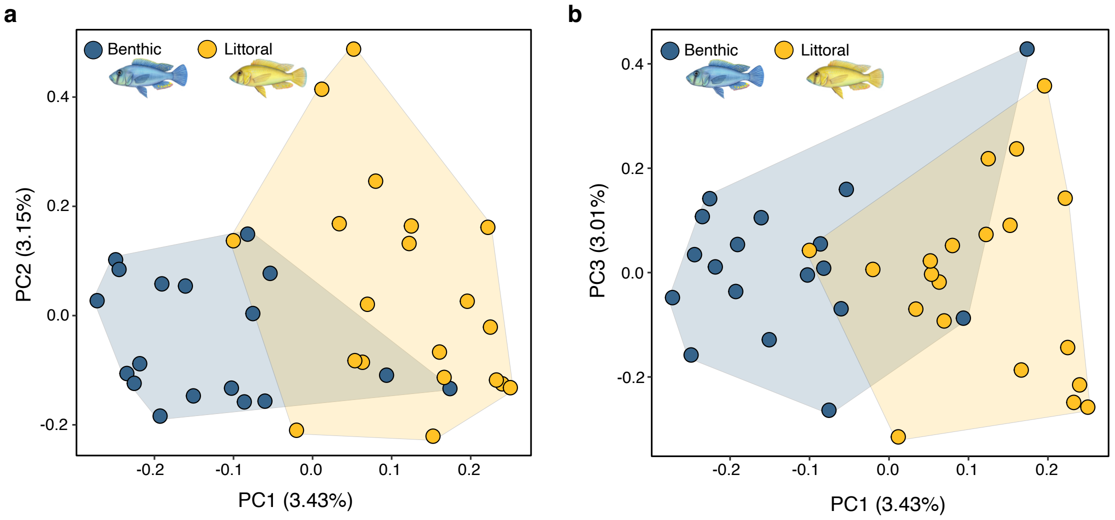


**Supplementary Figure 7 | Principal component plots of individual genotypes.** PCAs shows variation in individuals genotypes (n=38 individuals) along the first and second principal components **(a)**, and the first and third principal components **(b),** based on set of 89,718 high-confidence SNPs and indels identified across the transcriptome.

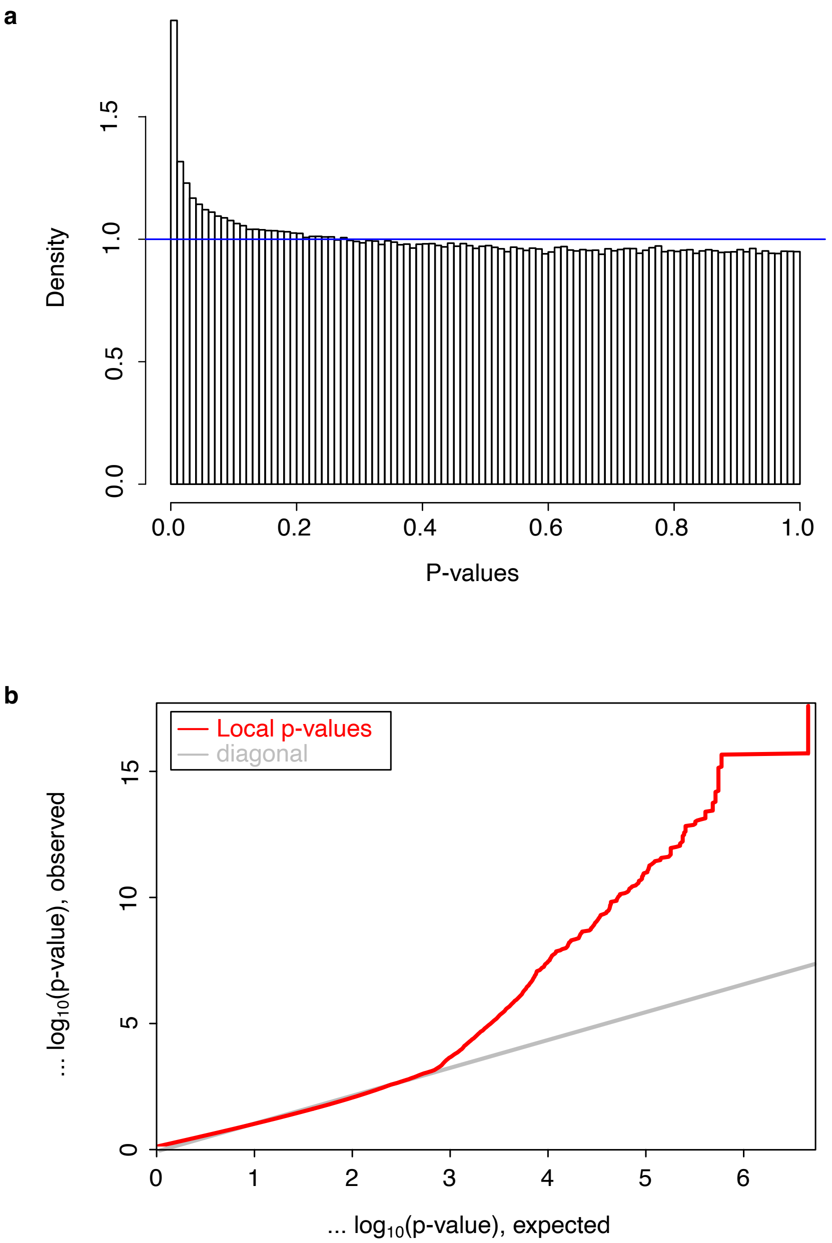


**Supplementary Figure 8 | Distribution of inferred p-values from eQTL analysis. a)** Histogram of p-values for all 3,615,374 *cis*-eQTLs tested with MatrixEQTL, showing no deviation that could have been caused by unaccounted covariates. **b)** Quantile-quantile plot distribution of p-values for all tested *cis*-eQTLs inferred by MatrixEQTL. The observed distribution of p-values deviates from the expected null distribution in the right tail, therefore spurious causes inflation was assessed using permutation tested of simulated datasets (see Supplementary Figure 9 and Supplementary Table 15). Both figures were obtained directly from MatrixEQTL.

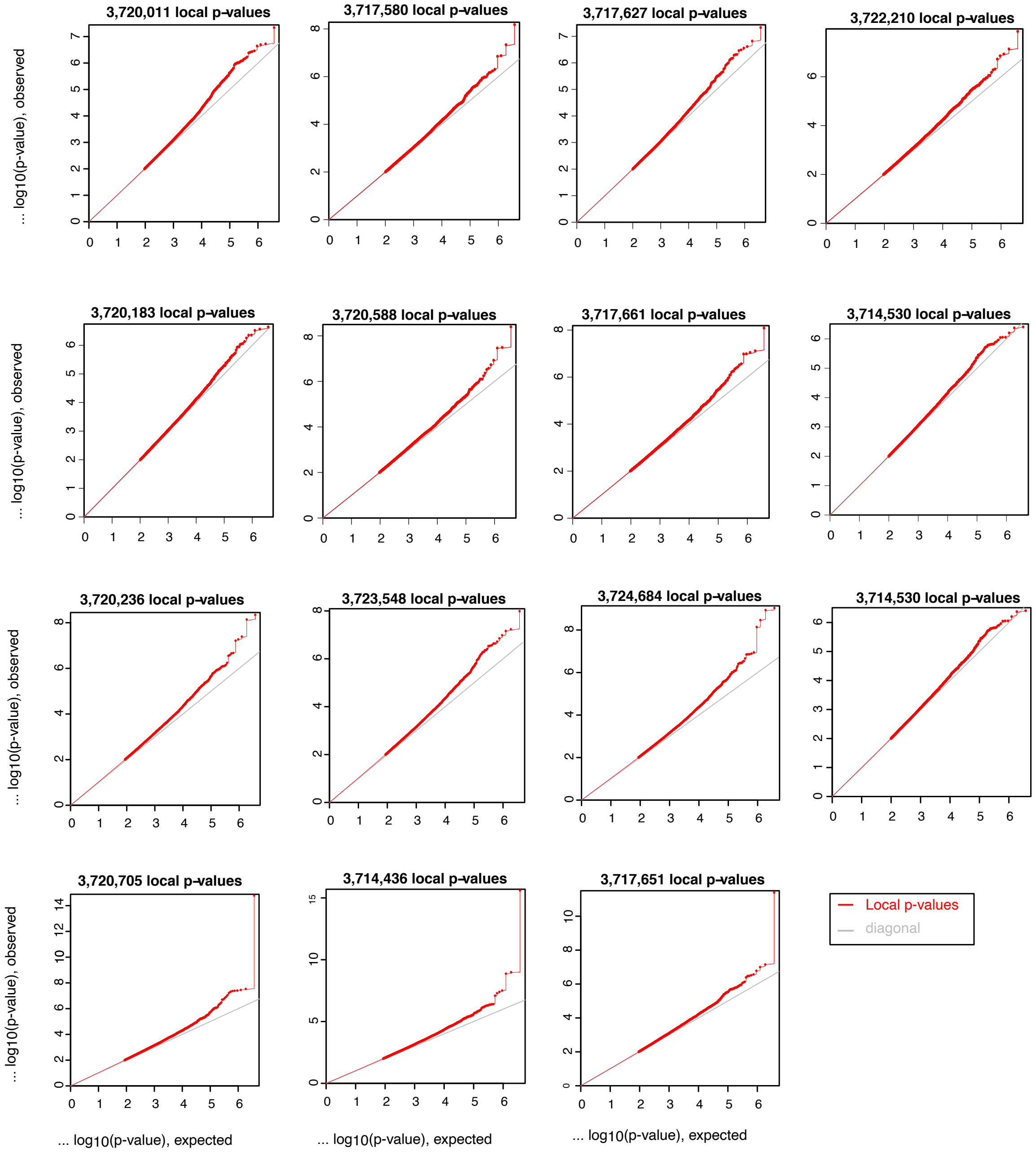


**Supplementary Figure 9 | Distribution of inferred p-values eQTL permutation tests.** Quantile-quantile plot distributions of p-values for *cis*-eQTLs generated for permutation simulated datasets permutation, showing the p-values generated for the first 15 *cis*-eQTLs simulated data models. The number of *cis*-eQTLs inferred by MatrixEQTL for each simulated model is given in the corresponding plot.

a

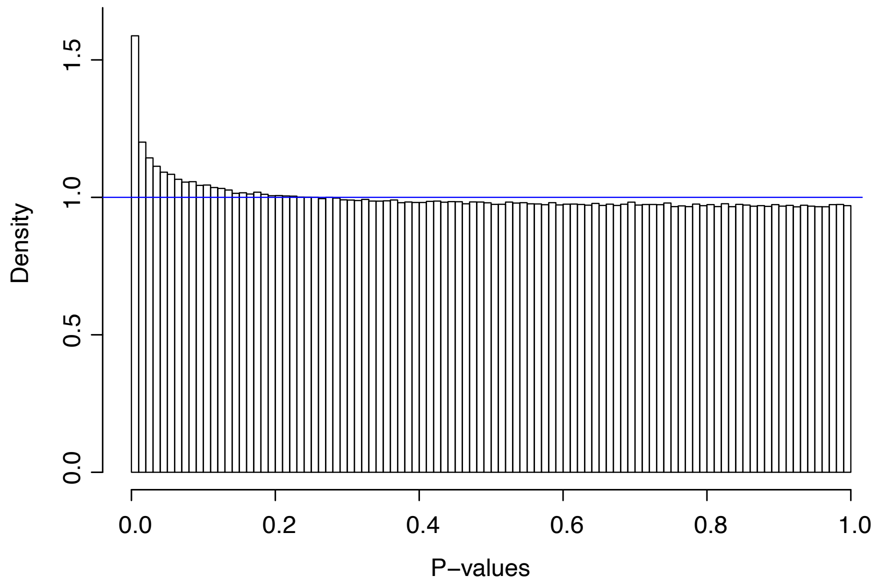


b

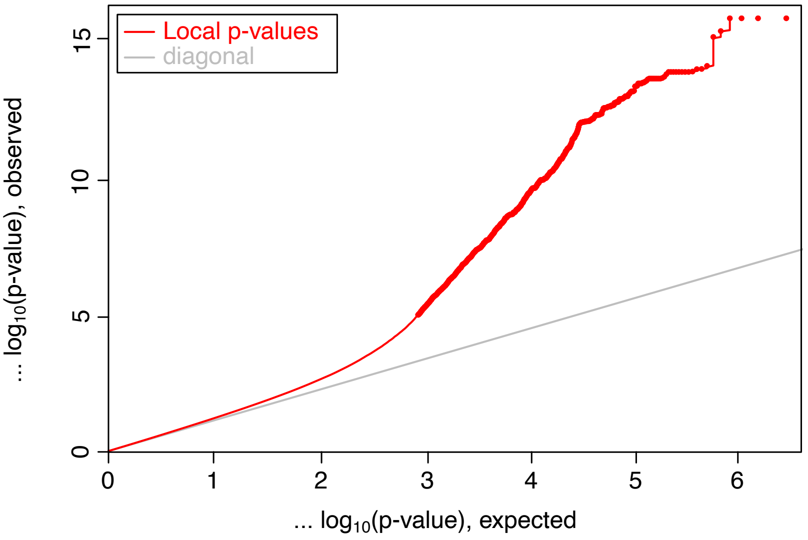


**Supplementary Figure 10 | Distribution of inferred p-values from sQTL analysis. a)** Histogram of p-values for all 17,170,992 *cis*-sQTLs tested with MatrixEQTL, showing no deviation associated with unaccounted covariates. **b)** Quantile-quantile plot distribution of p-values for all tested *cis*-sQTLs inferred by MatrixEQTL. The observed distribution of p-values deviates from the expected null distribution in the right tail, therefore spurious causes inflation was assessed using permutation tested of simulated datasets (see Supplementary Figure 11 and Supplementary Table 16). Both figures were obtained directly from MatrixEQTL.

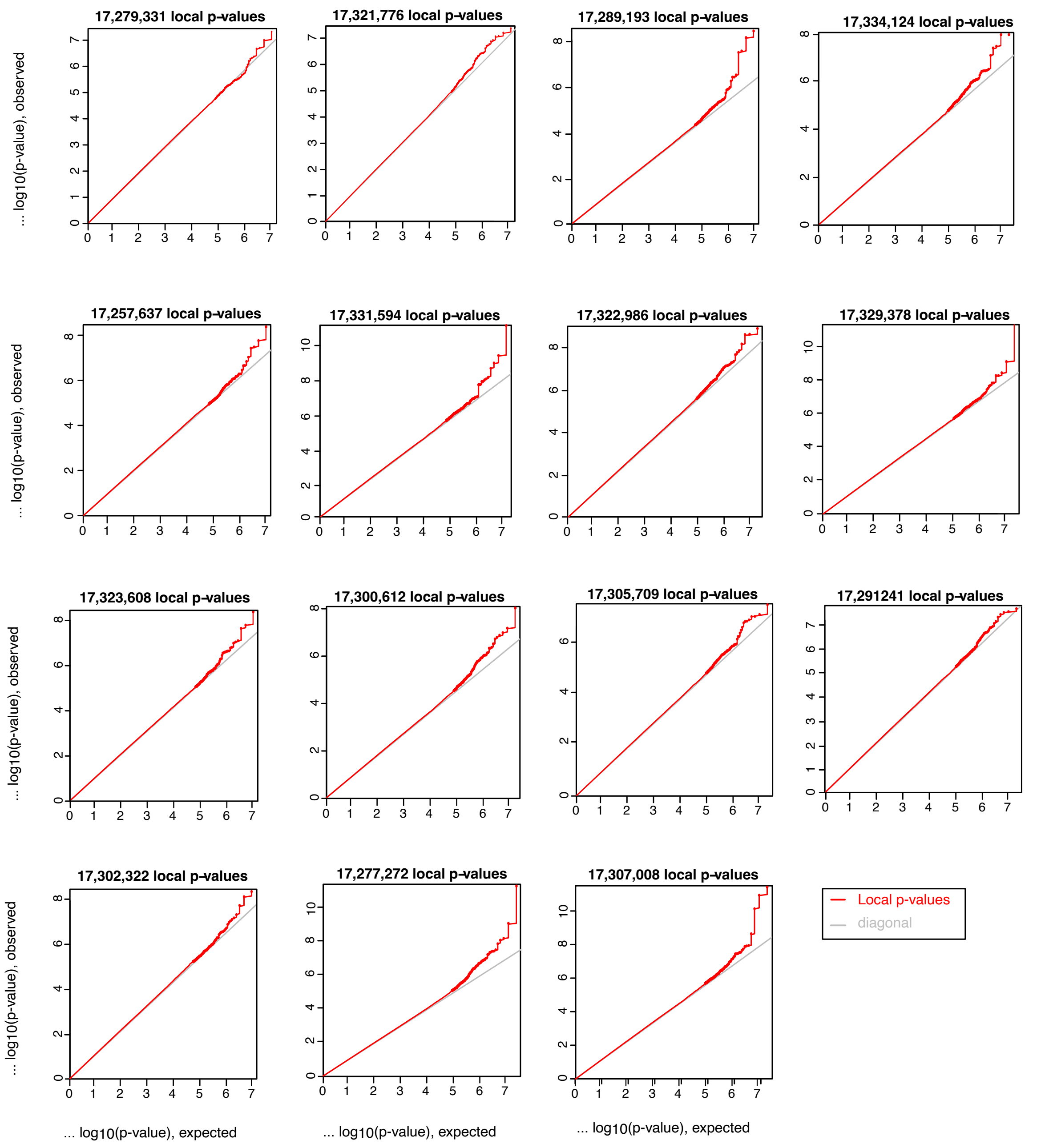


**Supplementary Figure 11 | Distribution of inferred p-values sQTL permutation tests.** Quantile-quantile plot distributions of p-values for *cis*-sQTLs generated for permutation simulated datasets permutation, showing the p-values generated for the first 15 *cis*-sQTLs simulated data models. The number of *cis*-sQTLs inferred by MatrixEQTL for each simulated model is given in the corresponding plot.

**Supplementary Tables**

All supplementary tables are provided in a single separate file.
